# Representation of verbal thought in motor cortex and implications for speech neuroprostheses

**DOI:** 10.1101/2024.10.04.616375

**Authors:** Erin M. Kunz, Benyamin Meschede-Krasa, Foram Kamdar, Donald Avansino, Samuel R. Nason-Tomaszewski, Nicholas S. Card, Brandon Jacques, Payton Bechefsky, Nick Hahn, Carrina Iacobacci, Leigh R. Hochberg, David M. Brandman, Sergey D. Stavisky, Nicholas AuYong, Chethan Pandarinath, Shaul Druckmann, Jaimie M. Henderson, Francis R. Willett

**Affiliations:** Department of Electrical Engineering, Stanford University, Stanford, CA, USA; Wu Tsai Neurosciences Institute, Stanford University, Stanford, CA, USA; Department of Neuroscience, Stanford University, Stanford, CA, USA; Department of Neurosurgery, Stanford University, Stanford, CA, USA; Howard Hughes Medical Institute at Stanford University, Stanford, CA, USA; Wallace H. Coulter Department of Biomedical Engineering, Emory University and Georgia Institute of Technology, Atlanta, GA, USA; Department of Neurological Surgery, University of California, Davis, CA, USA; School of Engineering and Carney Institute for Brain Sciences, Brown University, Providence, RI, USA; VA RR&D Center for Neurorestoration and Neurotechnology, Rehabilitation R&D Service, Providence VA Medical Center, Providence, RI, USA; Center for Neurotechnology and Neurorecovery, Department of Neurology, Massachusetts General Hospital, Harvard Medical School, Boston, MA, USA; Department of Neurosurgery, Emory University, Atlanta, GA, USA; Department of Cell Biology, Emory University, Atlanta, GA, USA; Department of Neurobiology, Stanford University, Stanford, CA, USA

## Abstract

Speech brain-computer interfaces show great promise in restoring communication for people who can no longer speak^1–3^, but have also raised privacy concerns regarding their potential to decode private verbal thought^4–6^. Using multi-unit recordings in three participants with dysarthria, we studied the representation of inner speech in the motor cortex. We found a robust neural encoding of inner speech, such that individual words and continuously imagined sentences could be decoded in real-time This neural representation was highly correlated with overt and perceived speech. We investigated the possibility of "eavesdropping" on private verbal thought, and demonstrated that verbal memory can be decoded during a non-speech task. Nevertheless, we found a neural "overtness" dimension that can help to avoid any unintentional decoding. Together, these results demonstrate the strong representation of verbal thought in the motor cortex, and highlight important design considerations and risks that must be addressed as speech neuroprostheses become more widespread.

## Introduction

For people with paralysis due to injury or disease, brain-computer interfaces (BCIs) offer a promising solution for restoring lost movement or communication function^7^. Successful demonstrations have shown that people with tetraplegia can use neural signals to control a computer cursor^8–13^ and to reach and grasp with a robotic arm, or even their own arm^14–16^. More recently, communication BCIs have restored rapid communication via accurate decoding of handwriting^17^ and speech^1,2,18–20^, substantially exceeding the communication rates offered by alternative devices (e.g., eye tracking). Encouragingly, the most recent demonstration has shown successful open-ended use of a speech neuroprosthesis for regular, everyday communication with friends, family and co-workers by a person with ALS^3^.

Given such rapid progress in speech BCIs, it is important to better understand the limits of what can be decoded from speech motor cortex. One concern by potential users is that of mental privacy, or whether a speech BCI would “be able to read into thoughts or internal monologues of users when attempting to decode (motor) speech intentions” ^6^. Most, but not all, people often associate conscious internal experiences of cognition with language, reporting that they have an “inner monologue” ^21–24^. Inner speech (also called imagined speech, internal speech, covert speech, silent speech, self-talk, speech imagery, internal monologue or verbal thought) is theorized to play an important role supporting complex cognitive processes including working memory, verbal rehearsal, logical reasoning, executive function, behavioral control and motivation^25–31^. Inner speech is also implicated in silent reading, with many people reporting evoked auditory or motor imagery of speech while reading ^32,33^.

Both neuroimaging and electrophysiological studies have shown that inner speech engages a similar (but not completely overlapping) cortical network as overt speech^34–36^, raising the possibility that electrodes placed in regions to decode overt speech may also enable inner speech decoding^34,37–47^. Precise neural differences between inner and overt speech remain largely elusive^25,32,48,49^. Neuroprosthetic studies using electrocorticography (ECoG) have begun to show evidence that inner speech can be decoded from particular regions of the cortex but differ in their conclusions about which regions contribute^46,50–52^. Most recently, Wandelt et al. demonstrated that inner speech could be decoded from signals recorded by intracortical microelectrode arrays in supramarginal gyrus (SMG), and that there was shared representation between words produced covertly, overtly, and heard^53^.

Here, we studied the neural representation of inner speech in three BrainGate2 speech BCI research participants with severe dysarthria using microelectrode arrays placed in the motor cortex. We discovered that inner speech is robustly represented, even in a task where the use of inner speech was not explicitly instructed, which raises the possibility of eavesdropping on private thought with a speech BCI. By characterizing its neural geometry, we found that inner speech appears to be a more weakly modulated version of overt speech. The neural similarity of inner speech and overt speech could cause a decoder to unintentionally decode inner thoughts. However, we also found that a large “overtness” neural dimension can help to distinguish between inner speech and overt speech, and that if speech BCIs are explicitly trained to ignore inner speech, they can do so with very high accuracy.

## Results

### Representation of inner speech in motor cortex

To investigate the representation of inner speech in the motor cortex, we analyzed microelectrode array recordings from three BrainGate2 clinical trial participants (T12, T15 and T16) during different types of verbal speech behaviors (Table 1). At the time of data collection, participants T12 and T15 were severely dysarthric due to amyotrophic lateral sclerosis (ALS) and participant T16 was dysarthric due to pontine stroke. All participants could partially articulate and vocalize, but were unable to produce speech intelligible to the untrained listener.

**Table 1:**
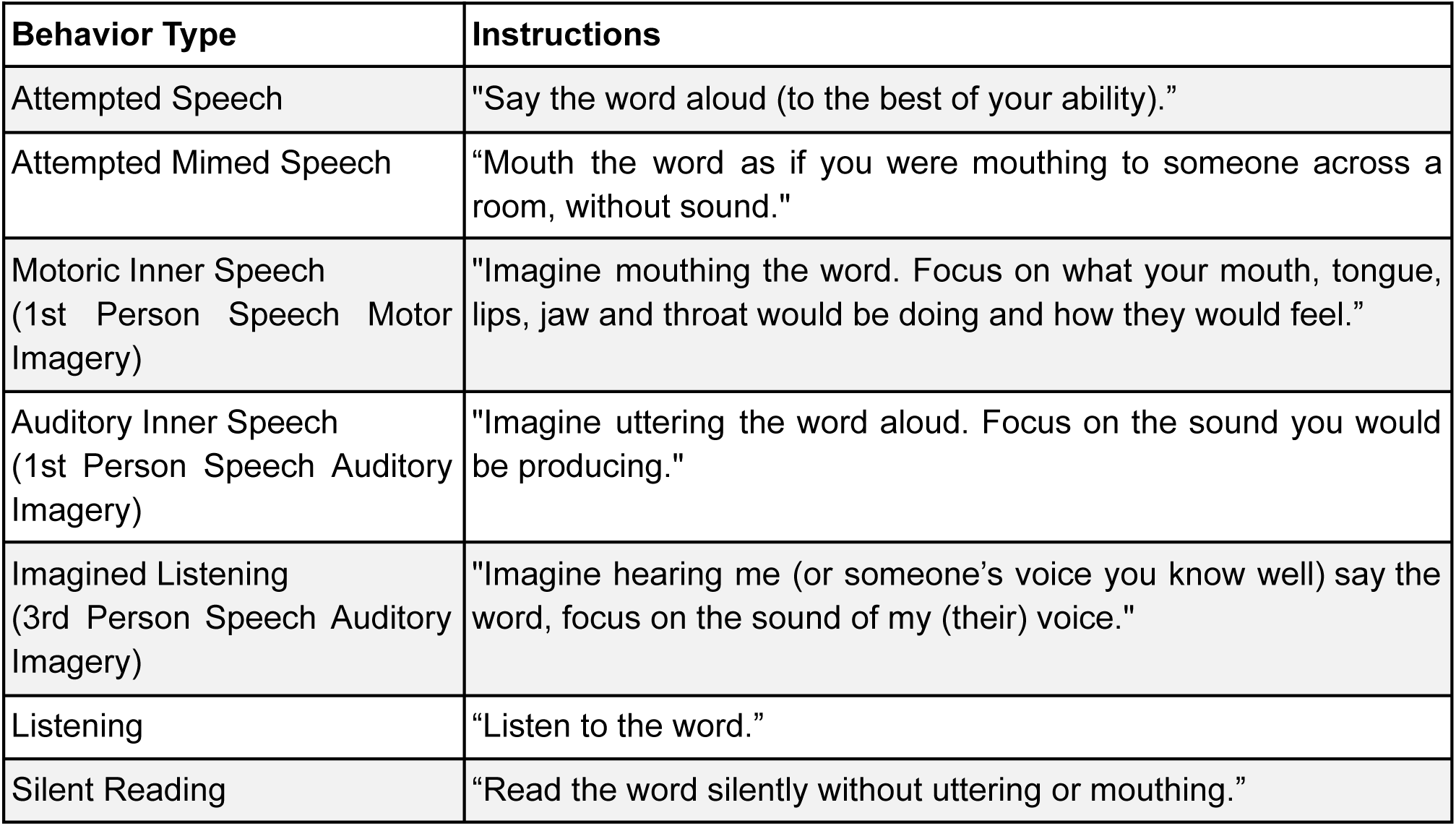
Task instructions for each verbal behavior condition.

The task instructions were chosen to explore the gradient between overt (attempted or mimed) and covert (inner) speech, perceived speech and silent reading. We cued three different strategies for covert speech (1st person auditory speech imagery, 1st person motoric speech imagery and 3rd person auditory imagery) based on the different ways in which people have reported experiencing inner speech ^25^.

Participants performed each behavior according to visual cues displayed on a computer screen in an instructed-delay task (Figure 1A-B). The word set included seven single-syllable English words that each consisted of a consonant-vowel-consonant phoneme sequence that shared no phonemes with any other word.

**Figure 1:**
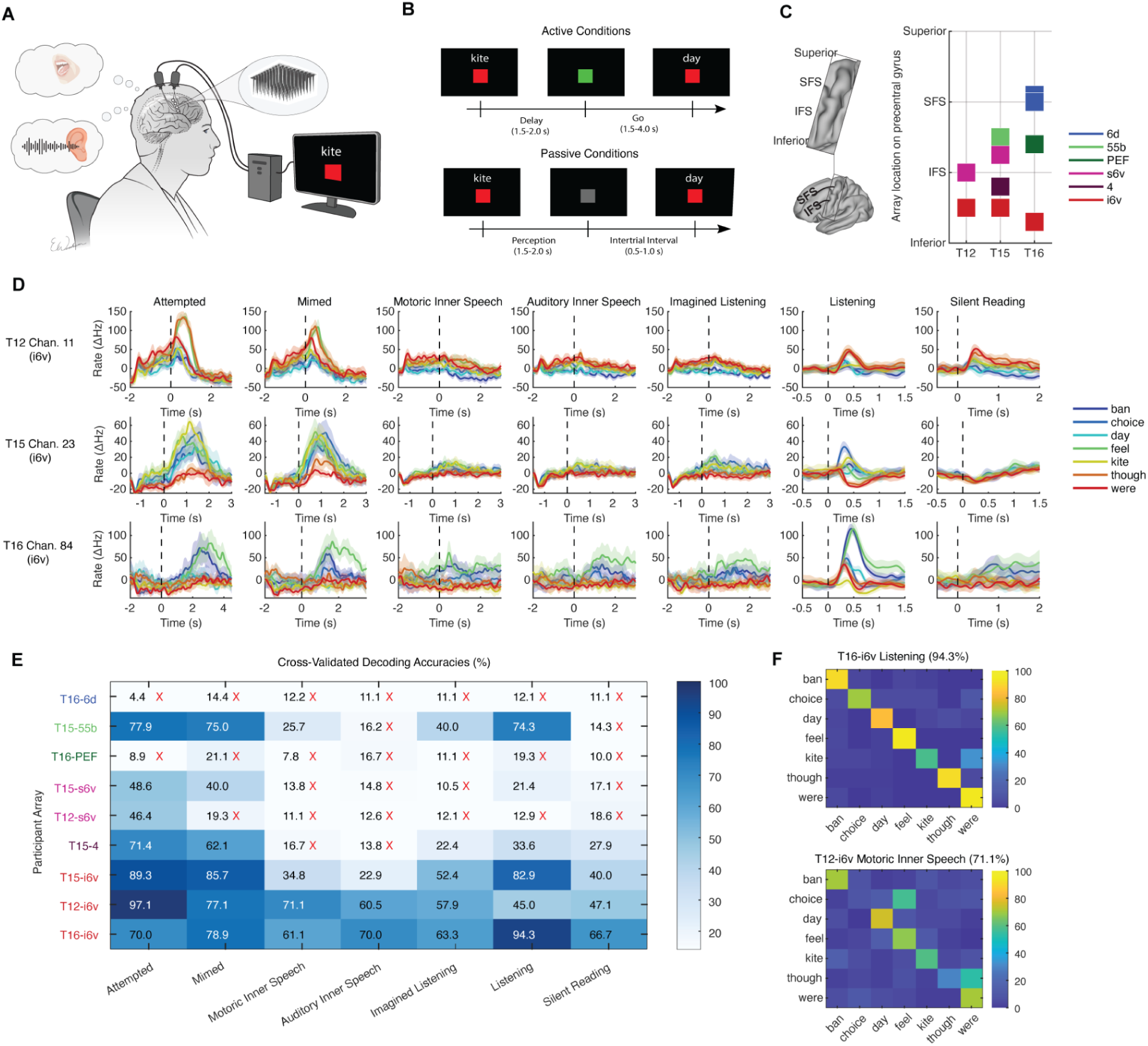
Inner speech, perceived speech and silent reading are represented in ventral and mid precentral gyrus. **A**. To assess tuning to different verbal behaviors, neural activity was recorded while participants were instructed to overtly or covertly produce speech, silently read or listen to speech. **B**. Example trial structure and visual cues shown on the screen for active (overtly or covertly producing speech) and passive (silent reading or listening) behavior conditions. No text was displayed during listening blocks. **C**. Neural activity was recorded from microelectrode arrays chronically implanted along the precentral gyrus in three participants. **D**. The mean firing rate recorded for each cued word for each behavior is shown for an example electrode from each participant. Shaded regions indicate 95% confidence intervals (CIs). Neural activity was smoothed by convolving with a Gaussian smoothing kernel (60-ms SD). **E**. Ten-fold cross-validated decoding accuracy by array (rows) and verbal behavior (columns) for a Gaussian Naive Bayes classifier applied to a 500 ms window of neural activity. Red X’s denote that the 95% confidence interval for decoding accuracy intersected chance level (14.3%). **F**. Example confusion matrices for Gaussian Naive Bayes classifiers trained on T16’s listening trials and T12’s motoric inner speech trials.

Participants had either 2 (T12) or 4 (T15, T16) multielectrode arrays placed along the precentral gyrus, spanning the functional regions of areas 6v, 4, PEF, 55b, and 6d as defined by individualized multi-modal cortical parcellations^54^ (Figure 1C). We analyzed array recordings from each functional region separately to more precisely understand localized representational differences.

We found that all the types of covert speech and perceived speech were represented in the precentral gyrus. More precisely, the arrays placed in the most inferior region of 6v (i6v) in all three participants had word representations that could be decoded above chance (14.3%) using a Gaussian Naive Bayes classifier for all seven behaviors (Figure 1E). T15’s 55b array (mid precentral gyrus) also had significant representation of two covert conditions and the listening condition. Surprisingly, in some participants the decoding accuracies for covert and perceived conditions approached or even exceeded that of overt conditions (T16’s listening could be decoded with 94.3% accuracy, compared to 70.0% and 78.9% for overt conditions, and T12’s motoric inner speech could be decoded with 71.1% accuracy, compared to 97.1% and 77.1% for overt conditions) (Figure 1F). The array in T15’s area 4 (primary motor cortex) showed weak representation of imagined listening, listening, and reading and T15’s superior 6v (s6v) array also showed weak representation of listening conditions. Other sampled regions either only had representation for the attempted condition (T12-s6v) or were in regions not considered part of the speech motor network, and appeared to show no speech representation (T16-6d ‘hand knob’ area, T16-PEF premotor eye field).

### A shared neural code for overt, covert and perceived speech

Having found representations for inner speech, listening and silent reading in the same regions of precentral gyrus, we next investigated the relationship between the neural representations of the same words across behaviors. We considered two hypotheses for how covert speech could be represented within the same neural ensemble as overt speech while not triggering any motor output. First, overt and covert speech could lie in orthogonal subspaces^55,56^, allowing for independent encoding of output-potent overt and output-null covert speech signals as has been found in motor cortex for reaching^57–59^. Alternatively, overt and covert speech could exist within the same neural space but differ in magnitude of modulation, such that covert speech does not reach an activation threshold to generate motor output ^60^.

We assessed overlap between the neural representations of overt and covert speech by measuring the correlation between firing rate vectors (where each electrode’s firing rate is an entry in the vector) evoked by the same word across different behaviors. Figure 2A shows an example correlation matrix generated by correlating firing rate vectors associated with each word and behavior. Strong off-diagonal banding indicates that words are encoded similarly across behaviors, including listening and silent reading. To summarize the strength of the correlations, we measured the average cross-behavior correlations across all arrays in regions that exhibited tuning to all behaviors, in all 3 participants (Figure 2B). Correlations were generally high across all tested behaviors, consistent with a behavior-agnostic neural representation of words.

**Figure 2:**
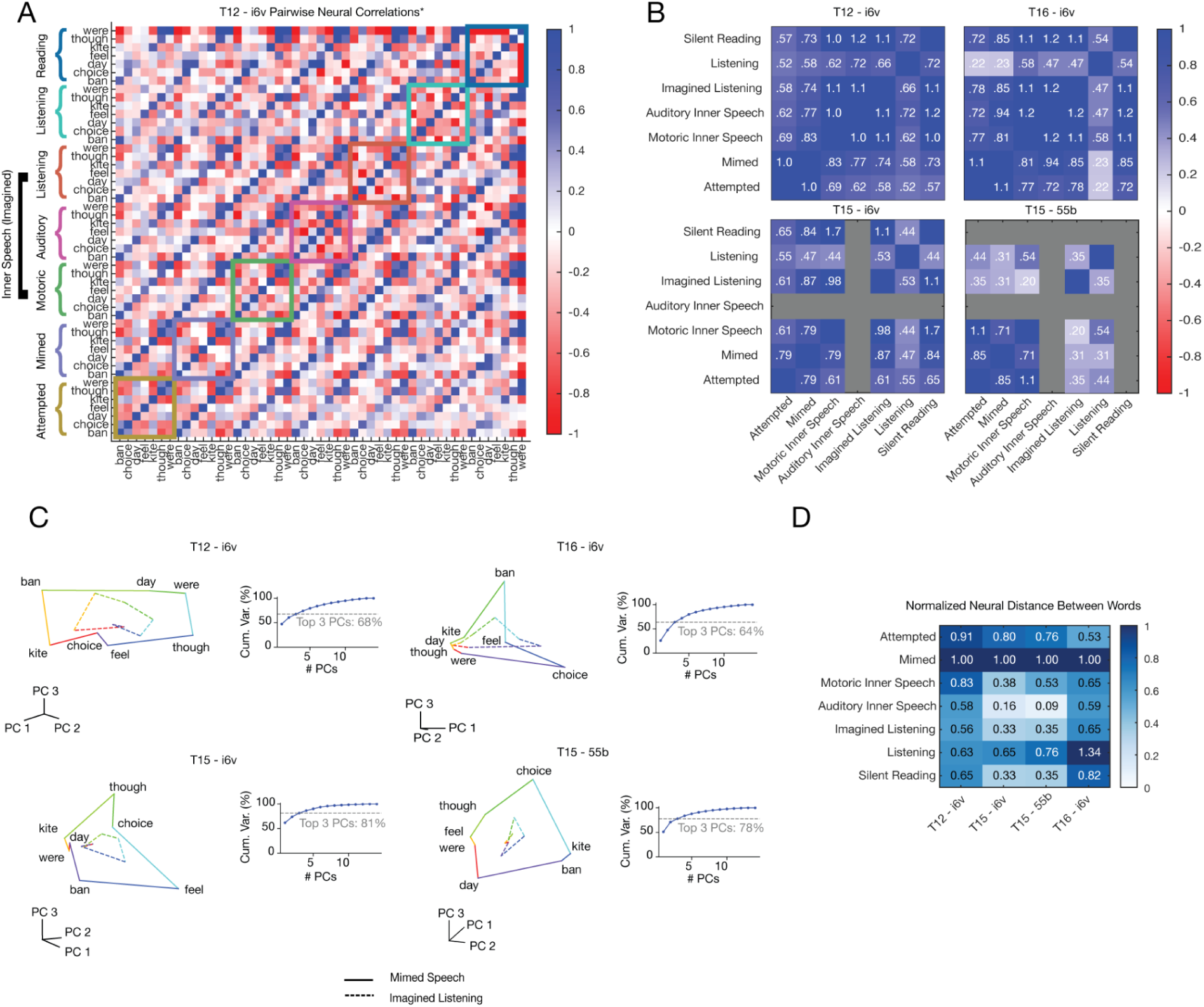
Inner speech and perceived speech as scaled-down versions of overt speech in motor cortex. **A** Each (i, j) entry of the correlation matrices indicates the Pearson correlation between the mean firing rate vector of word-behavior *i* and that of word-behavior *j*. The off-diagonal banding shows that the same word vectors across the different verbal behaviors were correlated. Correlation was computed using a cross-validated metric that reduces bias. **B**. Each (i, j) entry represents the average correlation of same-word pairs between behavior i and j. Since a cross-validated estimator of correlation was used, values can be greater than 1 (see methods for details). **C**. Projections of average word representations into the subspace defined by the top three principal components visually demonstrate the shared structure and relative sizes of word representations across behaviors. **D**. Average neural distances between words within each behavior, normalized to the attempted mimed behavior (largest). This metric represents the relative sizes of the word representations in neural population state space (i.e., for T12 imagined listening word representations are about 56% the size of mimed speech).

We used Principal Components Analysis (PCA) (see Methods) to visualize the neural geometry of the seven words in a three-dimensional space (Figure 2C), focusing on the overt mimed behavior and the covert imagined listening condition (since these behaviors were most strongly represented on average, across all 3 participants). We see that the relative positions of words is preserved across the behaviors, with the imagined listening condition “ring” appearing as a scaled down version of the mimed behavior “ring.” To quantify the relative size of the representations, we examined the neural distances between each pair of words, as a measure of the size of the neural representations (larger distances indicate greater separation between words due to greater magnitude of modulation). We normalized the distances by the most separable overt behavior (Attempted Mimed). The results indicate that while the structure of the seven word conditions across behaviors is highly correlated, the inner speech and perceived behavior representations are scaled down versions of the overt conditions (Figure 2D). The strong correlations across all behaviors for three participants in i6v indicate that this area of precentral gyrus contains representation of words that is largely behavior agnostic. This finding, in combination with the relative sizes of the behaviors, appears more consistent with the activation threshold hypothesis for how motor output is prevented during listening, reading and covert speech.

### Real-time decoding of continuous inner speech in three dysarthric individuals

We next investigated whether self-paced inner speech of whole sentences is also represented in motor cortex. To test this, we evaluated the performance of an inner speech BCI that decodes continuously imagined sentences. Such an approach could be preferred by some BCI users, since attempted speech requires activation of orofacial muscles and breath control that can be fatiguing and produces undesirable, distracting vocalizations.

Using the decoding pipeline from our previous work^1,3^ (Figure 3A), we instructed participants to covertly speak sentences using their preferred “inner speech” strategy and trained real-time decoders using this data. Approximately 1 hour of inner speech sentence data, as well as previously collected overt speech data, were used to train the decoders (see Methods for details). We then evaluated the online performance of the inner speech decoder with a set of 50 sentences derived from a 50-word vocabulary^1,18^. The inner speech online brain-to-text decoder achieved a 24%, 14% and 33% Word Error Rate for participants T12, T15 and T16 respectively (Figure 3B-D). To our knowledge, this is the first demonstration of an online, continuous inner speech decoder in severely dysarthric participants. All participants in this study were familiar with attempting to use overt speech in the brain-to-text framework and reported a preference for using inner speech instead, citing the lower physical effort required to perform inner speech, as well as the improved outward appearance of a neutral posture as compared to the strained, unintelligible vocalization accompanying attempted speech.

**Figure 3:**
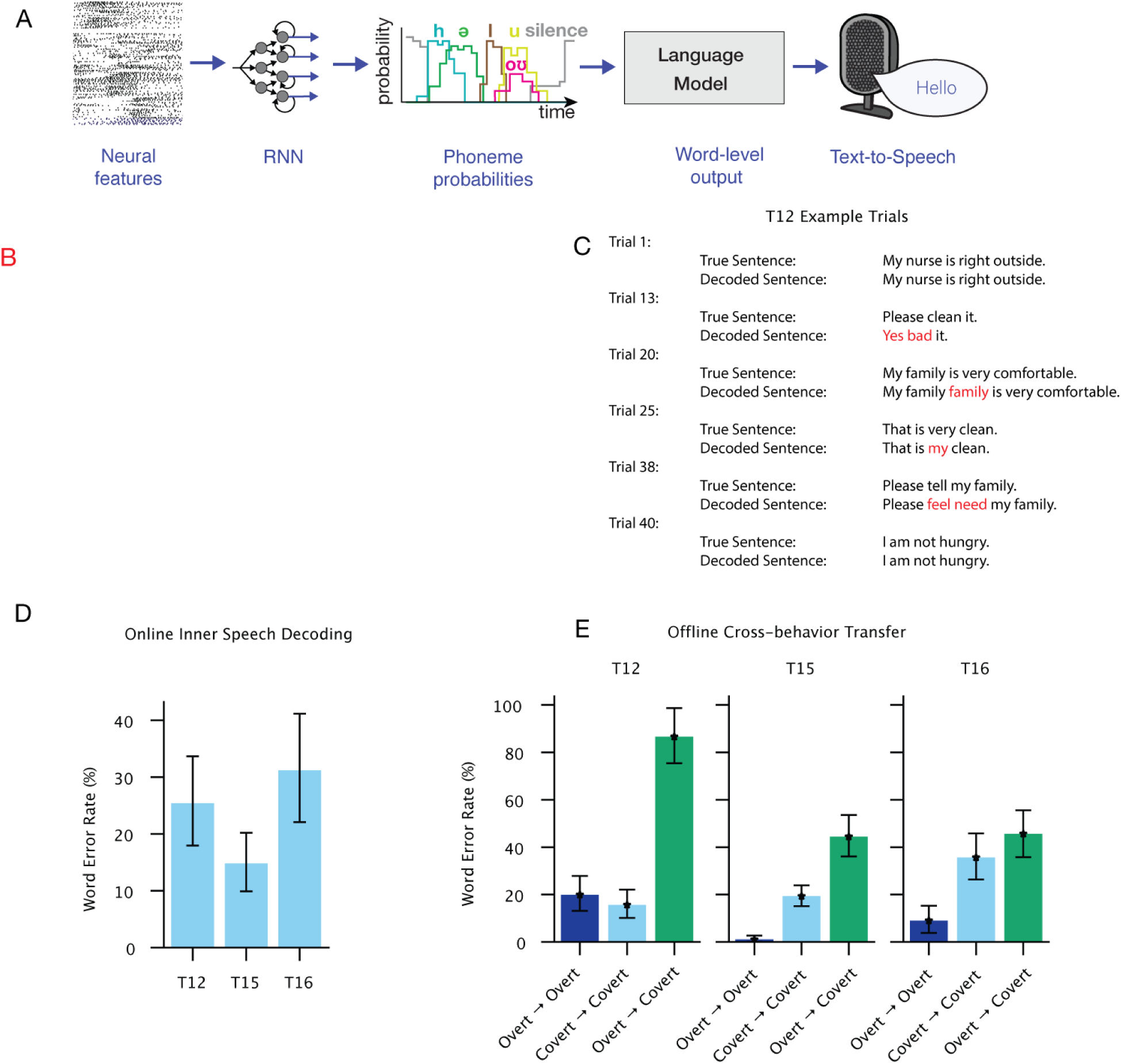
Real-time decoding of continuous inner speech. **A.** To decode neural features into text in real-time, neural features were passed into a recurrent neural network (RNN) which was trained to output probabilities for each of 39 phonemes and a silent token (SIL) at every 80 ms time step. The phoneme probabilities were then passed through a language model decoder to find the most likely sequence of words consistent with both the phoneme probabilities and language statistics. The decoded text was presented on the screen and articulated by a text-to-speech algorithm. **B**. T12 using an inner speech BCI in real-time. Cued text appears above the green square and decoded text lies below. **C**. Example decoded sentences from participant T12 from an evaluation block with an overall word error rate of 24%. **D**. Word error rates during online inner speech decoding for all three participants. **E**. Offline performance of a decoder trained on overt speech and evaluated on covert speech trials (green bars), compared to baseline decoding performance for overt speech (dark blue) and covert speech (light blue). Decoding performance was above chance in all conditions (tested with bootstrap resampling, see Methods), meaning that an overt speech decoder can decode covert speech better than chance.

Lastly, to further probe the relationship between overt speech and inner speech, and to determine if inner speech could be inadvertently decoded by an overt decoder, we collected the same training sentence set for both conditions (overt and covert). We trained decoders offline on these datasets and evaluated on test sets both within and across behaviors (Figure 3E). We found that, mostly as expected, overt speech offline decoding accuracies were better (T15 and T16, 95% confidence intervals were not intersecting) or on par (T12, 95% confidence intervals intersected) with their inner speech counterparts. Interestingly, when evaluating on inner speech test sentences with a decoder trained only on overt training data, we found performance was better than chance for all participants. This means that a brain-to-text decoder trained purely on overt speech signals from the motor cortex is capable of decoding inner speech.

### Uncued inner speech can can be decoded from ventral motor cortex

In all experimental tasks thus far, we investigated explicitly-cued inner speech; however, it is unknown to what extent the neural representation of uncued, private verbal thought correlates with the volitional inner speech behaviors reported above. To explore the potential to decode inner speech when it is not explicitly cued, a suite of sequential recall tasks were conducted with participant T12 in order to naturally elicit the use of inner speech. Previous studies on inner speech for cognition suggest that the nature of a cue can impact whether inner speech is used^61^ and that verbal short-term memory is a common strategy for maintaining sequentially presented information ^62^. Three upper extremity motor tasks were designed to variably elicit inner speech based on the nature of the visual cue (Figure 4A). No explicit instruction for mental strategy was given regarding whether or not to use inner speech to reproduce the cued instructions.

**Figure 4:**
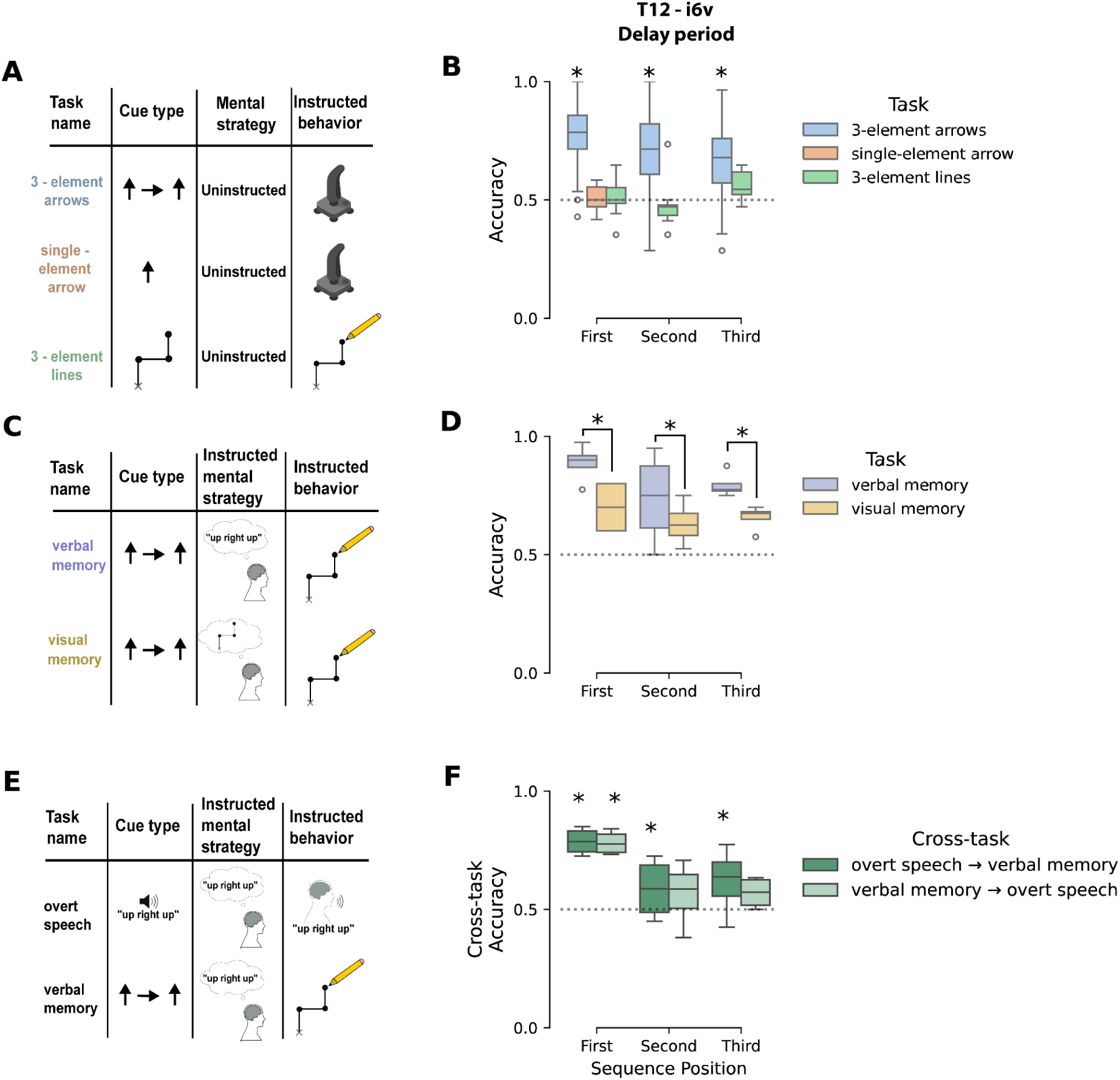
Uncued inner speech elicited by a serial recall task can be decoded from i6v. **A)** Participant T12 completed three upper extremity motor tasks designed with different cues and short-term memory requirements to variably elicit the use of verbal short-term memory. Mental strategy was not explicitly instructed. The 3-element arrows task was designed to elicit verbal short-term memory for serial recall, while the single-element arrow and 3-element lines tasks served as controls (and were designed *not* to elicit inner speech). **B)** Decodability of sequence positions was assessed by training binary Linear Discriminant Analysis models to classify between pairs of sequences that differ in a single position (e.g., first position: ↑ → ↑ vs ↓ → ↑). Models were fit to i6v neural activity using a 2 second window before the go cue. Cross-validated accuracy is plotted with chance performance indicated by the dotted black line. Box plots show the distribution of decoding accuracy across all possible pairs that differ only in one sequence position (10-fold cross-validated accuracy). Only the 3-element arrow elicited a significant neural representation of the sequence elements in all three positions during the delay period. Significance assessed via bootstrap-derived confidence intervals for decoding accuracy compared to chance level of 0.5 **C)** Participant T12 completed two versions of the same serial recall task, with explicit instruction for either a verbal or visual short-term memory mental strategy. Mental strategy refers to the researcher’s instructions to the participant on how to memorize the cued sequence. Instructed behavior refers to the instructed upper-extremity motor output during recall. **D)** Same as **B** but for tasks that only differed in instructed mental strategy. Decoding accuracy significantly increased in all sequence positions when T12 engaged in a verbal memory strategy. Significance assessed via bootstrap-derived confidence intervals of increase in decoding accuracy compared to chance level of zero **E)** An analogous overt speaking task was compared to an upper extremity motor task with instructed verbal strategy for short-term memory. **F)** Same as **B** except decoders are trained on overt speech (verbal memory) and tested on verbal memory (overt speech).

In the first task (Figure 4A ,“3-element arrows”), T12 was instructed to memorize a sequence of three arrows during the delay period, and then to sequentially move a joystick in those corresponding directions during the go period. We hypothesized that the symbolic nature of the arrow cues and the 3-element sequence length would elicit inner speech as a mental strategy to remember the sequence (e.g., by repeating “up right up, up right up, …” in an inner voice to remember the cue *↑* → **↑**). To quantify whether different sequence elements were represented during the delay period, binary decoders were trained to classify between cues differing in only one sequence position (e.g., above-chance performance for *↑* → **↑** vs ↓ → ↑ indicates that the first element is represented). In area i6v, all positions could be decoded above chance, potentially due to T12’s use of inner speech (Figure 4B; decoding accuracy confidence intervals (CIs) did not intersect the chance level of 0.5). Next, we tested two additional tasks designed *not* to elicit inner speech, which address the confound of a potential neural code for general movement sequencing in i6v (as opposed to a representation of inner speech specifically).

First, we tested a task variant in which only a single arrow cue was shown (Figure 4A, “single-element arrow”); due to its simplicity we hypothesized that T12 would not be inclined to use inner speech to accomplish the task. The same decoding analysis showed that in i6v, the cued direction was *not* decodable above chance in the single-element arrow task (Figure 4B, decoding accuracy CI did not intersect the chance level of 0.5). This demonstrates that the i6v tuning observed above for 3-element sequences was not simply due to motor planning or individual symbolic cues. Finally, a third task (3-element lines) was designed to elicit sequential movement of the hand without inner speech. Instead of using arrow symbols, a sequence of line segments were presented as an image, and T12 was instructed to reproduce the image by drawing the lines in an ordered sequence. We hypothesized that the geometric nature of the cue was less likely to elicit inner speech than the arrow cues in the first task. Consistent with this hypothesis, movement directions were not decoded significantly above chance for any sequence position in the line task (Figure 4B, decoding accuracy CIs did not intersect the chance level of 0.5). Taken together, these tasks show that the representation of movement sequences seen in the 3-element arrows task is unlikely to be caused by tuning to hand-motor preparation or planning of sequences in general, and is instead consistent with uncued inner speech.

To further test our hypothesis that 3-element arrow cues elicited the unprompted use of inner speech, we asked T12 to perform the 3-element arrows cued motor sequence task, while explicitly being instructed to either use verbal or visual memory for sequence memorization (Figure 4C). T12 performed the task with 100% accuracy for correct sequence recall (Supp. Figure 2) and reported being able to reliably switch between strategies. In i6v, tuning significantly increased in all sequence positions when T12 volitionally used a verbal memory strategy (Figure 4D, CI for increase in decoding accuracy did not cross chance level of 0).

Finally, to probe whether the representation of neural activity during instructed verbal memory was similar to attempted speech (which would be expected due to their shared neural representation shown in Figure 2), T12 performed two sequence recall tasks. The first was cued with audio of spoken direction sequences and T12 was instructed to attempt to speak the direction sequence (Figure 4E, overt speech). The second was identical to the verbal memory task in figure 4C. The same sequence-position decoding analysis was performed, except now with decoders trained on one task and evaluated on the other. We found that decoders could generalize across tasks significantly above chance in all three positions (Figure 4F, decoding accuracy Cis did not intersect the chance level of 0.5). This indicates that i6v representations of inner speech used for verbal short-term memory are encoded similarly to overt speech. Taken together, these controls confirm that the delay-period decodability of the 3-element movement sequences seen above (Figure 4B) was most likely due to T12’s uninstructed use of verbal short-term memory, and demonstrates that a decoder trained on overt speech can be used to decode a private, cognitive use of inner speech which was never intended to be vocalized aloud.

We attempted to replicate these results with participant T16 and found similar results, though the effect was weaker and only existed during the go period (Supp. Figure 1). This could suggest that there is individual variation in the encoding strength and timing during which verbal short term memory is decodable from i6v.

### Evidence of a robust behavior dimension separating overt and covert speech

The shared neural code for covert and overt speech, as well as the decodability of uninstructed verbal short term memory during simple tasks, raises the risk of unintentional inner speech decoding. However, in our prior experiment which showed high neural correlations between overt and covert behaviors (Figure 1,2), behaviors were tested in separate experimental “blocks.” This means that two behaviors collected in separate blocks could appear to have erroneous differences in neural activity due to spurious drifts in firing rates known to occur across time^63,64^. Because of this, we could not conclude if there were large differences in mean firing rate across behaviors that could help a decoder to distinguish between overt and covert speech. To test for this, we conducted a follow-up experiment in which both overt and covert speech trials of the seven words were randomly interleaved within the same experimental blocks, making it possible to conclusively test for differences in mean firing rates across behaviors.

First, we visualized the relative structure of word conditions in the top three principal components of PCA fit on all 14 of the interleaved conditions (7 words each for overt and covert speech, Figure 5A-C). The top three components captured between 64% and 82% of the variance, depending on the participant. The low-dimensional representation again reveals that the neural representations of inner speech share the same relative structure between words as the overt speech condition while being a scaled down version, as expected from prior results above. However, rotating the same projection also reveals strong separability between the two behaviors along an “overtness dimension” which expresses a large mean-shift between covert and overt speech.

**Figure 5:**
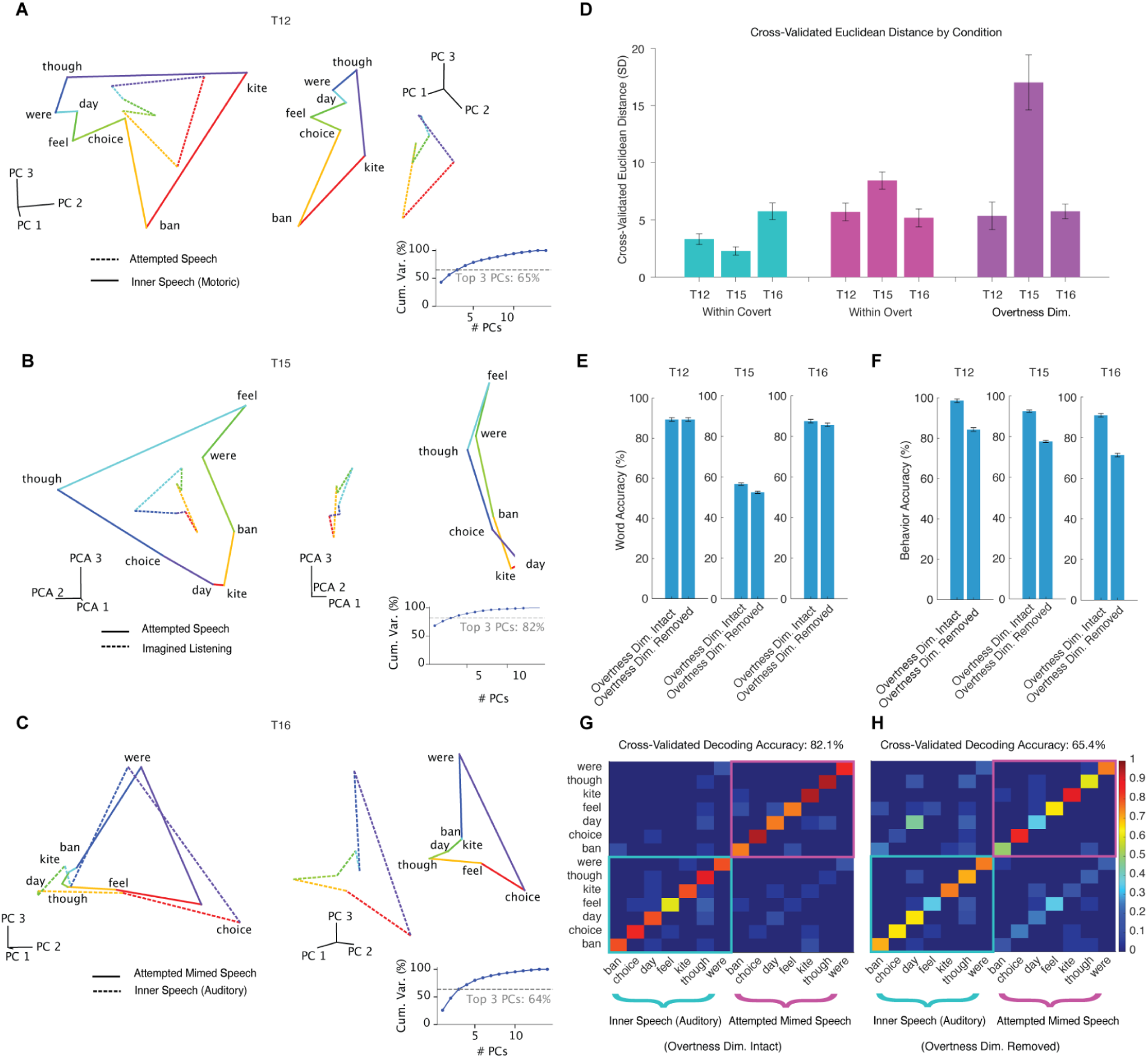
Motor cortex contains a neural dimension representing overtness that can help distinguish covert speech from overt speech. **A**. Projections of average word representations into the subspace defined by the top three components of Principal Components Analysis (capturing 65% of variance) on all 14 conditions demonstrate shared structure between behaviors (left), but also strong separation between the word “rings” when axes are rotated to visualize the overtness dimension that captures differences in mean rates between behaviors (right). **B**. PCA projections for T15. **C**. PCA projections for T16. **D**. Cross-validated Euclidean distances between word pairs within each behavior (turquoise and pink), representing the size of word-related neural modulation , and distances between matching word pairs across behaviors (purple), representing the relative magnitude of the separating overtness dimension. The overtness dimension contains behavior-modulation similar in size (T12, T16) or larger than (T15) word-related modulation. **E**. Word accuracy (indicating correct word decoding irrespective of the decoded behavior) before and after within-behavior mean subtraction (removal of the overtness dimension). Removing the overtness dimension does not substantially affect word decoding accuracy. **F**. Behavior accuracy (indicating correct behavior decoding irrespective of the decoded word) before and after within-behavior mean subtraction (removal of the overtness dimensions); behavior-decoding is adversely affected for all 3 participants **G,H.** Exemplary confusion matrices for T16 of the classifier across all 14 conditions before (G) and after (H) the overtness dimension was removed.

To better understand the size of the overtness-related modulation relative to word-related modulation, we compared the Euclidean distances in neural state space between word pairs within each behavior (word-related modulation) to distances between matching word pairs across behaviors (overtness modulation) (Figure 5D). While the size of word-related modulation varied across participants and behaviors, overtness modulation was comparable to (T12, T16) or greater than (T15) word-related modulation, indicating that it is a robust feature of the neural activity that could help decoders to distinguish between covert and overt speech.

To understand the potential effect of the overtness dimension on decoding, we trained Gaussian Naive Bayes decoders to distinguish between all 14 conditions, both with the overtness dimension intact and with the overtness dimension removed. We removed the overtness dimension by mean-subtracting each neural feature across all trials within each behavior. Decoder performance with the overtness dimension intact showed little confusion across the behaviors, while the decoder trained on data without the overtness dimension showed increased off-diagonal confusion of like-words across behaviors, as expected (Figure 5G,H show T16 results). However, the decoding performance remained relatively high, potentially due to the differing sizes of the word representations between the behaviors that could be used as additional information by the decoder. To confirm that the signal being removed indeed contained predominantly behavior-specific information (as opposed to word-related information), we calculated the word and behavior accuracies before and after removing the overtness dimension (Figure 5E,F). As expected, word accuracy (whether the correct word was predicted even if the behavior was confused) remained similar for both conditions, but behavior accuracy (whether the correct behavior was predicted even if the word was confused) decreased substantially for all 3 participants.

### RNN decoders can be trained to ignore inner speech with very high accuracy

Having found that certain neural dimensions specifically encode behavior, and that simple classifiers can distinguish between covert and overt speech, we next investigated whether RNN decoders could be trained to distinguish between the two in the more applicable, continuous sentence decoding context. To do this, we explored an additional “imagery-silenced” decoder training strategy in which we trained an RNN using overt training sentences (which were labeled normally with their constituent phonemes) combined with inner speech training sentences (which were labeled instead with a “silence” token) (Figure 6A,B). When tested offline, this imagery-silenced strategy largely preserved decoding performance on the overt behavior (Figure 6C) while robustly preventing content from being decoded during the inner speech trials (Figure 6D). To further understand the effect of this imagery-silenced technique compared to its “imagery-naive” counterpart (i.e., an RNN trained only on overt trials), we analyzed the logit outputs of both RNNs for each behavior and found that logits from matching overt and covert sentences were highly correlated in the imagery-naive case, but were much less correlated in the imagery-silenced case (Figure 6E). Examining the output phoneme probability logits from individual trials highlights that the imagery-silenced strategy dramatically quiets RNN output on inner speech trials (Figure 6F), although this strategy remains to be verified in online, real-time decoding.

**Figure 6:**
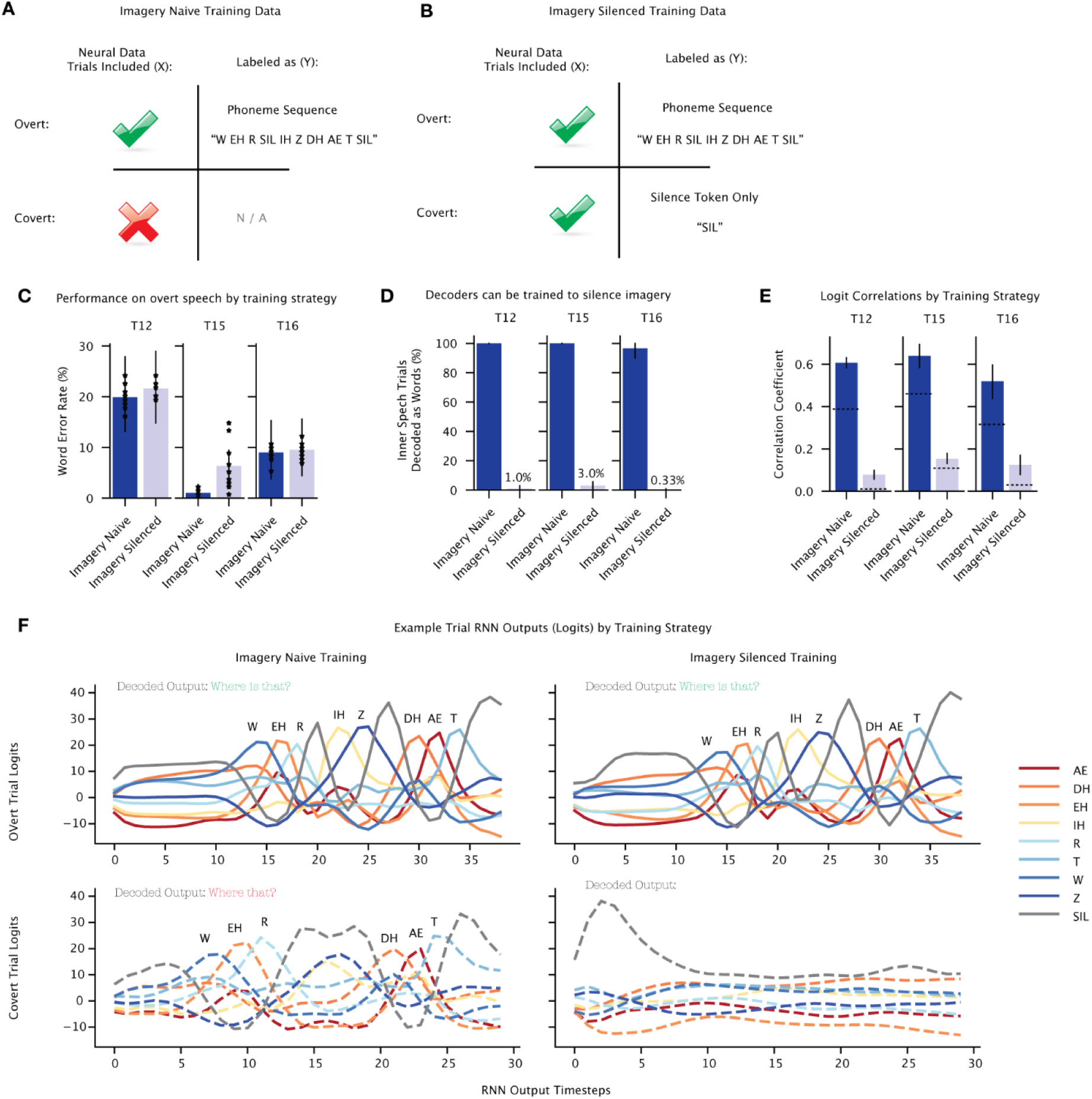
An imagery-silenced training strategy for RNNs can robustly prevent accidental covert speech decoding. **A**. The current training strategy for most real-time speech BCIs is limited to overt speech trials only. **B**. We tested an imagery-silenced training strategy in which we also included covert speech sentence trials that were labeled as silence (i.e. the silent phoneme token). **C**. The imagery-silenced training strategy largely preserved offline decoding performance (word error rate) on overt speech trials for each participant (dots indicate 10 RNN training seeds, error lines are 95% confidence intervals). **D**. Imagery-silenced training robustly prevents false outputs during covert speech. **E**. Correlations between the RNN outputs for matched covert and overt speech sentences are much higher in the imagery-naive training regime than in the imagery-silenced regime. Dotted line indicates the chance-level correlation for that scenario (see Methods). **F.** Visualization of phoneme logit outputs for one T16 sentence for the overt trial (top) and covert trial (bottom) from a decoder trained with the imagery-naive strategy (left) versus the imagery-silenced strategy (right).

## Discussion

The recent progress in speech neuroprostheses has raised questions regarding the extent to which private verbal thought may be accessible via neural recordings from speech-related areas of motor cortex. Here, we found that decodable representations of inner speech, perceived speech and silent reading exist in localized regions of the motor cortex (middle and most ventral region of precentral gyrus). Furthermore, these representations are highly correlated with overt speech, raising the possibility that an overt speech BCI may inadvertently decode inner or perceived speech. We further demonstrate that a speech BCI could potentially “eavesdrop” on private, uncued verbal thought by showing that inner speech can be decoded during tasks which naturally elicit the use of verbal short term memory. However, further investigation also identified a neural dimension that encodes “overtness,” and we demonstrated that when explicitly trained to do so, decoders can be robustly prevented from decoding inner speech unintentionally.

We also demonstrated, for the first time, an online continuous inner speech neuroprosthesis that decoded sentences from a 50 word vocabulary with a word error rate between 14-33% across 3 participants. Due to improved ease-of-use, outward appearance and a potential increase in speed, an inner speech neuroprosthesis could be preferable to a BCI based on attempted speech, assuming that volitional intent to communicate with inner speech can be reliably distinguished from inner speech that is meant to be kept private.

### Verbal representations in motor cortex

Speech is a complex, high-dimensional behavior unique to humans that has been found to engage various regions of the cortex in a “speech network”. Differing from traditional views that posit separate speech perception and speech production areas ^65–67^, recent studies have shown that the neural mechanisms of speech production and perception are connected and overlapping, even in motor areas^68–71^. Studies on inner speech have also shown overlapping mechanisms between overtly and covertly produced speech ^34,46,50–53^. However, these studies disagree on the degree of overlap and role of the speech motor cortex during covert and overt speech production, and speech perception.

With the level of spatial resolution achieved with microelectrode array recording in three participants, this work demonstrates that there are localized regions of the motor cortex - the middle (55b) and most ventral (i6v) regions of precentral gyrus - for which perceived, overt and covert speech and silent reading share a similar neural representation. The regions lying in between these (4, s6v) and more dorsally (6d) showed neural tuning to overt production but not perception or covert production, or lacked strong neural representation of speech altogether.

This suggests that the neural substrates in areas 55b and i6v have an abstract representation of speech that goes beyond the low-level control of motor articulators, and that is relatively agnostic to the verbal behavior. This aligns with previous speech BCI studies in which the middle precentral gyrus^2^ and the most ventral part of precentral gyrus^1,3^ demonstrated the greatest signal contributions to speech decoding.

The fact that these “speech hotspot” regions (areas 55b and i6v) had strong representations for all verbal behaviors raises the question of how motor output is inhibited during inner speech. One possibility is that overt and covert speech could lie in orthogonal neural subspaces^55,56^, allowing for independent encoding of output-potent overt and output-null covert speech signals, similar to what has been found in motor cortex for the preparation and execution of reaching ^57–59^. However, we found that the neural representations of overt and covert speech and even listening were highly correlated and thus largely in shared neural space. These shared signals could represent an abstract sensorimotor or auditory “goal” or target signal used by downstream motor areas, where output gating could occur instead. Alternatively, the weaker modulation evoked by inner speech in areas 55b and i6v may simply not be strong enough to cross an activation threshold required to generate motor output^60^; or, aspects of the neural activity unique to overt behavior (such as the “overtness” dimension we found here) could be used to gate motor output, as has recently been suggested for imagined wrist movement in hand motor cortex^72^.

### Implications for Intracortical Speech BCIs

In this work, we demonstrated the first real-time covert speech BCI in three people with severe dysarthria. Using a covert speech-based BCI proved to be favorable compared to one based on overt speech in terms of the lower physical effort required and improved outward appearance. Additionally, covert speech lacks the physiological constraints that slow down attempted overt speech in dysarthric individuals (e.g. difficulty controlling breathing while speaking) and could offer a path forward for the communication rate of speech BCIs to reach that of typical conversation rates. A covert speech BCI may require additional design considerations to prevent accidental “leakage” of inner thought into BCI output, which might be accomplished by a volitional intent neural decoder, or an on/off switch (that could be operated with e.g. eye tracking, or a neurally decoded “click”).

Beyond potential design improvements to speech BCIs, this work also highlights ethical risks of decoding speech from speech-motor cortex. One major concern is the risk of unintentional decoding of inner speech or private thought by a BCI trained only to recognize overt speech, a concern which falls under the category of “executory user control” described in^6^. We show that despite the shared neural code for overt and covert speech and listening, careful design choices during decoder algorithm training can safeguard against unintentionally decoding inner speech. Additionally, ^3^ report that speech decoding systems that record from precentral gyrus can detect attempted speech with high specificity even without such design considerations. On the other hand, we demonstrate that for verbal short-term memory during a non-speech sequential recall task, the contents of a covert mental strategy for sequence memorization can be decoded. Inner speech as a cognitive tool in adults has also been implicated in task switching, planning, propositional reasoning, reasoning about others, spatial orientation, categorization, cognitive control, and reading^25^. Whether inner speech in these contexts is decodable from area i6v or 55b remains unknown. Furthermore, the degree to which speech and language are used for thought is under debate^73,74^. The scope of the risk of unintentional decoding of thought may be limited to concrete mental strategies such as verbal mental repetition for short-term memory. Private inner monologue likely differs between individuals and may not unravel concretely, which could make it difficult or impossible to decode from motor cortex.

Another risk for end users of speech BCIs is adversarial decoding of inner speech not intended to be externalized or shared. We define adversarial decoding as any verbal thought that is decoded from a speech BCI user’s neural data without their permission or knowledge. We demonstrate that algorithms can be trained on overt speech and leveraged to decode verbal short-term memory, listening, and perhaps other forms of private thought. This highlights the importance of data privacy for neural signals, particularly those recorded intracranially, which can contain decodable inner speech, and supports calls for strong privacy protections of neural data and informed consent for users considering speech BCIs ^75,76^. On the other hand, we show that covert speech is less strongly encoded relative to overt speech, thus limiting the ability to reliably decode it even with the highest resolution recording technology currently available. These metrics should be re-evaluated as new recording devices are used for speech decoding which may increase the feasibility of adversarially decoding verbal thought.

## Acknowledgements

We thank participants T12, T15, and T16 and their caregivers for their generously volunteered time and dedicated contributions to this research; B. Davis, K. Tsou and S. Kosasih for administrative support.

Support was provided by an ALS Pilot Clinical Trial Award (AL220043) from the Office of the Assistant Secretary of Defense for Health Affairs, a New Innovator Award (NIH 1DP2DC021055) from the National Institutes of Health and managed by the National Institute on Deafness and Other Communication Disorders, a grant (872146SPI) from the Simons Collaboration for the Global Brain, a postdoctoral fellowship funded by the A.P. Giannini Foundation, the Office of Research and Development, Rehabilitation R&D Service, Department of Veterans Affairs (nos. N2864C and A2295R), Wu Tsai Neurosciences Institute, Howard Hughes Medical Institute, Larry and Pamela Garlick, Simons Foundation Collaboration on the Global Brain and NIDCD (nos. U01-DC017844 and U01-DC019430), a Ketterer-Vorwald Neurosciences Interdisciplinary Graduate Fellowship, National Institute of Neurological Disorders and Stroke (NIH DP2NS127291), and a postdoc fellowship funded by the Eunice Kennedy Shriver National Institute of Child Health and Human Development.

*The contents do not represent the views of the Department of Veterans Affairs or the US Government. CAUTION: Investigational Device. Limited by Federal Law to Investigational Use.

## Declaration of Interests

The MGH Translational Research Center has a clinical research support agreement (CRSA) with Axoft, Neuralink, Neurobionics, Precision Neuro, Synchron, and Reach Neuro, for which LRH provides consultative input. LRH is a co-investigator on an NIH SBIR grant with Paradromics, and is a non-compensated member of the Board of Directors of a nonprofit assistive communication device technology foundation (Speak Your Mind Foundation). Mass General Brigham (MGB) is convening the Implantable Brain-Computer Interface Collaborative Community (iBCI-CC); charitable gift agreements to MGB, including those received to date from Paradromics, Synchron, Precision Neuro, Neuralink, and Blackrock Neurotech, support the iBCI-CC, for which LRH provides effort.

SDS is an inventor on intellectual property licensed by Stanford University to Blackrock Neurotech and Neuralink Corp. He is currently an advisor to ALVI Labs and is a member of the advisory board for Sonera. He is also a shareholder in Wispr.ai.

CP is a consultant for Meta (Reality Labs) and Synchron. DMB is a surgical consultant for Paradromics Inc.

SDS and DMB are inventors of intellectual property related to neuroprostheses owned by the University of California, Davis.

SD is a consultant for CTRL Labs.

JMH is a consultant for Neuralink and Paradromics, serves on the Medical Advisory Board of Enspire DBS and is a shareholder in Maplight Therapeutics. He is also the co-founder of Re-EmergeDBS. He is also an inventor on intellectual property licensed by Stanford University to Blackrock Neurotech and Neuralink Corp.

FRW is an inventor on intellectual property licensed by Stanford University to Blackrock Neurotech and Neuralink Corp.

All other authors have no competing interests.

## 1 Experimental Methods

### 1.1 Study Participants

Data from three participants, referred to as T12, T15, and T16, are reported in this study, all of whom gave informed consent and were enrolled in the Braingate2 Neural Interface System clinical trial (<u>ClinicalTrials.gov</u> Identifier: NCT00912041, registered June 3, 2009). Approval for this pilot clinical trial was granted under an Investigational Device Exemption (IDE) by the US Food and Drug Administration (Investigational Device Exemption #G090003), as well as the Institutional Review Boards of Stanford University (protocol #52060), University of California Davis, and Emory University (protocol #STUDY00003070). T12 gave consent to publish photographs and videos containing her likeness. All relevant guidelines and regulations were strictly upheld.

T12, a left-handed woman, was 68 years old at the time of data collection, with slowly-progressive bulbar-onset Amyotrophic Lateral Sclerosis (ALS) diagnosed at age 59 (ALS-FRS score of 26 at the time of study enrollment). On March 30, 2022, four 64-channel, 1.5 mm-length silicon micro electrode arrays coated with sputtered iridium oxide (Blackrock Microsystems, Salt lake City, UT) were implanted in T12’s left hemisphere, based on preoperative anatomical and functional magnetic resonance imaging (MRI) and individualized Human Connectome Project (HCP) cortical parcellation (see ^1^ for details). Two arrays were placed in HCP-identified area 6v (orofacial motor cortex) of ventral precentral gyrus, and two were placed in HCP-identified area 44 of inferior frontal gyrus (considered part of Broca’s area). Data are reported from post-implant days 412-742 . At the time of data collection, T12 was severely dysarthric for nearly 8 years due to bulbar ALS. She retained partial use of her limbs, and communicated primarily through use of a writing board or iPad tablet. She was able to vocalize while attempting to speak, and was able to produce some subjectively differentiable vowel sounds. However, we had difficulty discerning nearly all consonants produced in isolation and could not reliably make out any consonants or vowels when T12 attempted to speak full sentences at a fluent rate.

T15, a left-handed man, was 45 years old at the time of data collection, with ALS diagnosed at the age of 40. On July 17, 2023, four 64-channel, 1.5 mm-length silicon micro electrode arrays coated with sputtered iridium oxide (Blackrock Microsystems, Salt lake City, UT) were implanted in T15’s left hemisphere, based on preoperative anatomical and functional magnetic resonance imaging (MRI) and HCP individualized cortical parcellation (see ^2^ for details). Two arrays were placed in HCP-identified area 6v (orofacial motor cortex) of ventral precentral gyrus, one in HCP-identified area 55b, and one in HCP-identified primary motor cortex (area 4). Data are reported from post-implant days 230-333. T15 had no functional use of his upper and lower extremities and had severe dysarthria (ALS-FRS score of 23 at the time of study enrollment).

T16, a right-handed woman, was 52 years of age at the time of this study, with tetraplegia and dysarthria due to a pontine stroke approximately 19 years prior to enrollment in the BrainGate2 clinical trial. On December 2023, T16 had four 64-channel intracortical microelectrode arrays (Blackrock Microsystems, Salt Lake City, UT; 1.5 mm electrode length) placed in her left precentral gyrus, guided by individualized HCP cortical parcellation: two in HCP-identified hand knob area (area 6d), one in HCP-identified ventral premotor cortex (6v), and one on the border of the HCP-identified premotor eye fields (PEF) and speech-related 55b. Implant targets were guided by a multimodal cortical parcellation ^3^ of the left precentral gyrus. T16 was able to speak slowly and quietly, but enunciation was restricted by limited face and mouth movement. She had limited voluntary control of her upper extremities, with some shoulder motion and some slow and contractured wrist and finger movements. She had limited to no voluntary control of her lower extremities. T16’s sensation was fully intact. Data are reported from post-implant days 88-223.

### 1.2 Functional MRI Speech Lateralization & Array Placement

Prior to surgery, all participants underwent anatomic and functional brain imaging for speech and language lateralization, surgical planning and array placement targeting. (see ^1,2,4^ for array location estimates and further details).

### 1.3 Neural signal processing

Voltage time series signals were recorded using the Neuroplex-E system (Blackrock Microsystems) and transmitted via a cable attached to a percutaneous connector. Signals were analog filtered (4th order Butterworth with corners at 0.3 Hz to 7.5 kHz), digitized at 30 kHz (250 nV resolution) and fed into custom MATLAB software for digital filtering and feature extraction. To isolate signals relevant for estimation of neural ensemble activity ^5^, voltage time series were digitally high-pass filtered (250Hz cutoff) non-causally on each electrode using either a 1ms (T15) or 4ms (T12, T16) delay, and linear regression referencing (LRR) ^6^ was applied. Electrode-specific thresholds and LRR filter coefficients were determined using data recorded from an initial diagnostic block at the beginning of each session (see 1.5 Instructed Delay Tasks).

Next, two estimates of neural ensemble activity were computed for each electrode in either 10ms or 20ms bins. Threshold crossings were computed by counting the number of times the filtered voltage time series crossed an amplitude threshold set at either -3.5 or -4.5 times the standard deviation of the voltage signal. Default parameters for bin size and threshold varied between individual clinical trial site defaults. Spike band power was computed by taking the sum of squared voltages observed during each time bin. Threshold crossing rates and spike band power are estimates of local spiking activity and have been shown to be comparable to sorted single unit activity in terms of decoding performance and neural population structure ^7–9^. For each block, within each electrode, mean threshold crossing rates and spike band power was subtracted from each sample to account for neural nonstationarities (drifts in mean firing rate) which could arise over the course of a session ^10,11^. Threshold crossings and spike band power features were also z-scored (divided by their standard deviation) for all analyses apart from the PSTHs (Figure 1D), to ensure that electrodes with high firing rates did not overly influence the population-level results.

### 1.4 Data collection rig

Digital signal processing and feature extraction were performed on a dedicated computer. For T12, Simulink Real-time was used for data processing and the Psychophysics Toolbox ^12^ in MATLAB was used to implement task software. An additional Windows computer controlled task starting and stopping and interfaced with the Neuroplex-E system. For T15 and T16, BRAND ^13^ was used to implement modular, Python-based neural data processing and task software.

### 1.5 Overview of Data Collection Sessions

**Table 1:**
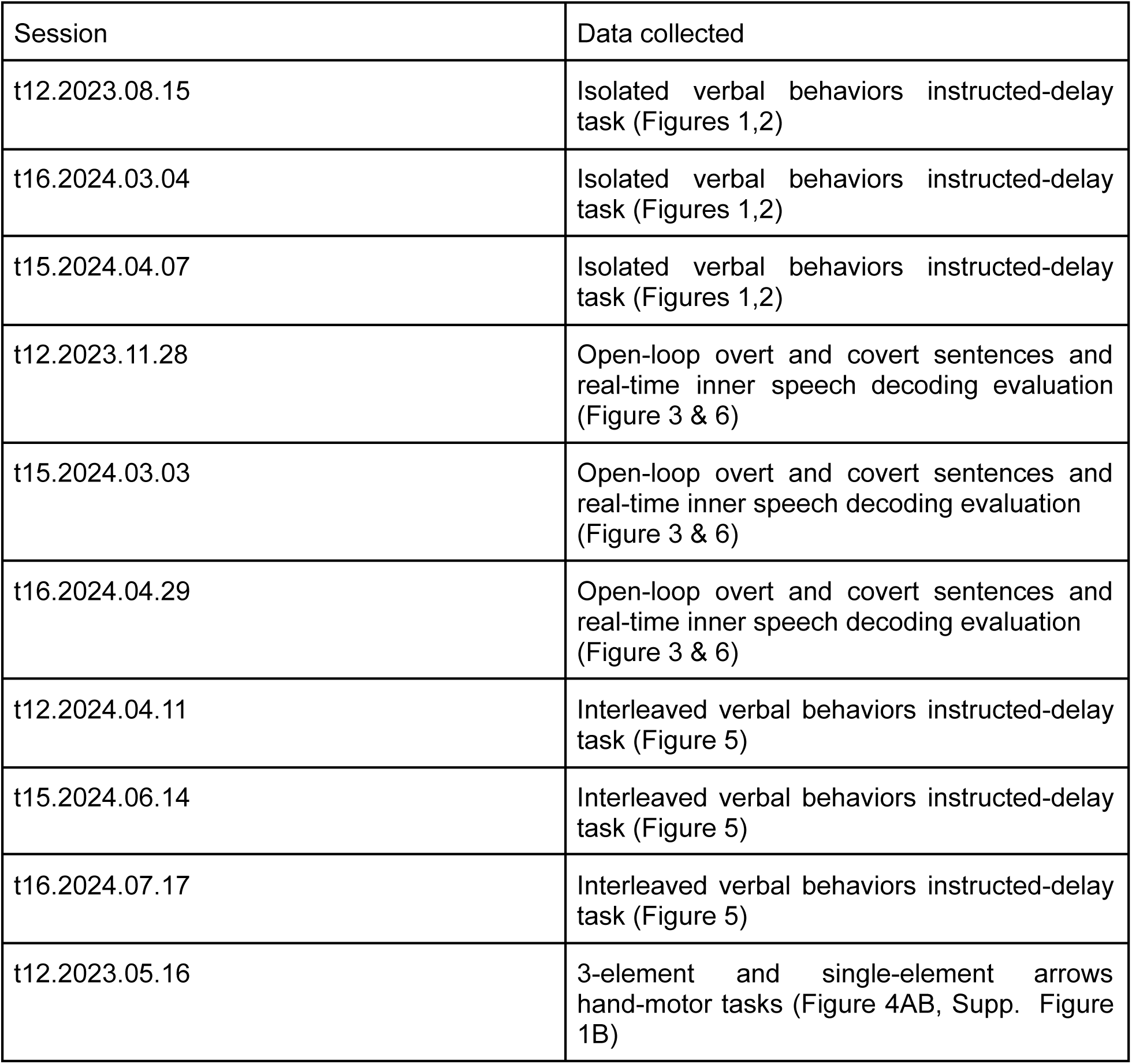
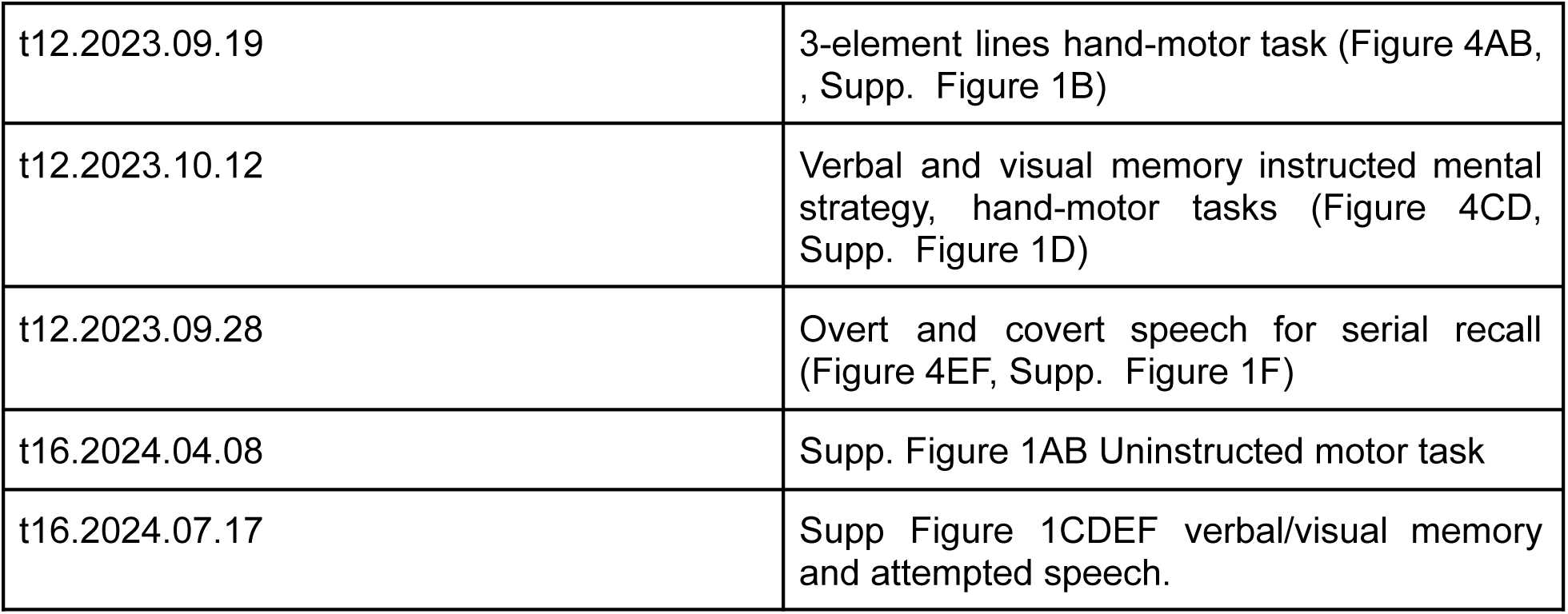
Data Collection Sessions.

## 2 Representation of Verbal Behaviors in motor cortex (Figure 1)

### 2.1 Isolated Verbal Behavior Instructed-Delay Task Details

For each research session that investigated ‘isolated’ verbal behavior (where each verbal behavior was tested separately in its own blocks), participants performed each verbal behavior in an instructed-delay task. The instructed-delay task for the “active” conditions, in which the participants either overtly or covertly spoke, consisted of a “delay” period and an execution, or “go,” period. The text cue was presented during the delay period with a red square signifying not to perform any behavior. Then, the square turned green while the text cue simultaneously disappeared at which point the participant began performing the desired verbal behavior. For the “passive” conditions (silent reading or listening) the trial design consisted simply of go-periods during which the text appeared (silent reading) or an audio file played a recording of the word being spoken (listening)., followed by intertrial intervals for the participant to return to baseline resting state in between trials. Trials were grouped into individual experimental “blocks” by behavior, and several blocks of each behavior were alternated throughout a research session. Exact durations of each trial period for a given verbal behavior were determined based on individual participants’ comfort and attention level. Timings and total number of trials by behavior are shown below.

**Table 2:** Delay/Intertrial Interval (ITI) Timing of instructed-delay verbal behavior tasks by participant.

|  | T12 | T15 | T16 |
| --- | --- | --- | --- |
| Attempted | 2s | 1.5s | 2s |
| Attempted Mimed | 2s | 1.5s | 2s |
| Motoric Inner Speech | 2s | 1.5s | 2s |
| Auditory Inner | 2s | 1.5s | 2s |
| Imagined Listening | 2s | 1.5s | 2s |
| Passive Listening | 1.5s (ITI) | 0.5s (ITI) | 1.5s (ITI) |
| Silent Reading | 1.5s (ITI) | 0.5s (ITI) | 1.5s (ITI) |

**Table 3:** Go Timing of instructed-delay verbal behavior tasks by participant.

|  | T12 | T15 | T16 |
| --- | --- | --- | --- |
| Attempted | 3s | 3s | 4s |
| Attempted Mimed | 3s | 3s | 3s |
| Motoric Inner Speech | 3s | 3s | 3s |
| Auditory Inner | 3s | 3s | 3s |
| Imagined Listening | 3s | 3s | 3s |
| Passive Listening | 1.5s | 1.5s | 1.5s |
| Silent Reading | 2s | 1.5s | 2s |

**Table 4:** Number of repetitions of each word collected of each verbal behavior task by participant.

|  | T12 | T15 | T16 |
| --- | --- | --- | --- |
| Attempted | 21 | 20 | 14 |
| Attempted Mimed | 21 | 20 | 14 |
| Motoric Inner Speech | 28 | 30 | 14 |
| Auditory Inner | 28 | 30 | 14 |
| Imagined Listening | 28 | 30 | 14 |
| Passive Listening | 21 | 20 | 21 |
| Silent Reading | 21 | 20 | 14 |

### 2.2 PSTHs (Figure 1D)

The time series shown in Figure 1D depict the average threshold crossing rates across time for each stimulus (word). 95% confidence intervals of the mean were computed by assuming that the data are normally distributed.

### 2.3 Naive Bayes classification (Figure 1E,F)

Offline classification results (reported in Fig 1d, Fig 3d,e) were generated using a cross-validated (leave-one-out) Gaussian Naive Bayes classifier, following the methods described in ^14^. Concatenated binned threshold crossing rates and spike band power were smoothed using a 60ms Gaussian kernel to reduce high frequency noise. Due to variations in the timing of neural modulation across behaviors and participants, we optimized the starting time of a 500 ms decoding window separately for each participant and behavior. In order to avoid biasing the decoding accuracies upwards due to overfitting the window start time, we used a nested 10-fold cross-validation strategy for window optimization. For each outer fold, the window start time was optimized on the training data using an inner 10-fold cross-validation to estimate decoding accuracy for each possible start time. The highest-performing start time was then chosen and applied to the test set of the outer fold, ensuring that decoding performance was never measured on data that was used to select the start time (note that this means decoding results could be aggregated across different windows, if the same start time was not chosen for each fold). A mean decoding accuracy was determined to be “significant” if the lower bound of the confidence interval was above the chance value of 14.3%. We chose to use a Gaussian Naive Bayes classifier because it is a simple method that performed well enough to demonstrate the existence of strong neural tuning - it is likely that more advanced methods could improve classification accuracy further.

## 3 Shared structure across verbal behaviors (Figure 2)

In Figure 2 we further analyzed the isolated verbal behavior task data. Binned threshold crossing rates and spike band power features were smoothed with a 60ms Gaussian kernel, and then averaged within a 500 ms window chosen for each behavior based on the optimization sweep done for Figure 1E (yielding a 128 x 1 feature vector per trial, for each 64 electrode array). Specifically, for each participant and behavior, the mode of the optimal window start times from the nested cross-validation analysis was chosen, as it should capture the most relevant word-specific modulation.

### 3.1 Cross-Validated Correlation Metric (Figure 2A,B)

Correlations (reported in Figure 2A) were estimated using a cross-validated method described in ^14^. Note that this bias-reduced estimator allows for a resulting value to be greater than 1 (or less than -1), particularly when the neural modulation is weak. Values greater than 1 (or less than -1) should be interpreted as evidence that the true correlation is near 1 (or -1). The average correlation of paired-words between behaviors is reported in Figure 2B. Note, we removed behaviors for which the neural modulation was too weak to accurately measure correlation (behaviors for which decoding accuracies were less than 25%).

### 3.2 Normalized Neural Distance (Figure 2C)

In Figure 2C, we estimated the average Euclidean distance in neural state space between all pairs of words within each behavioral condition separately, using the cross-validated distance estimator described in ^14^. Within each participant, we normalized the distances by dividing by the average distance of the “largest” overt behavior. In all participants, this was the attempted mimed condition.

### 3.3 Principal Components Analysis Visualization (Figure 2D)

To better visualize the neural geometry of covert and overt speech, we plotted projections of the data in the top 3 principal components as determined by Principal Components Analysis (PCA). First, for each word and behavior, neural activity was time-averaged and averaged across trials, yielding a 128 x 1 representation vector (for each 64-electrode array). These vectors were stacked to create a 128 x 14 matrix (7 words x 2 behaviors), and PCA was then applied across the columns of this matrix. We then plotted each of the 7 words within the 3-dimensional space created by the top 3 principal components, connected nearby words with lines to form "word rings" to better visualize the relative positions of each word (lines were drawn and colored consistently across behaviors), and rotated the viewpoint to best reveal the relationship between the two word rings shown.

## 4 Online inner speech decoder

### 4.1 Session Design

Real-time continuous inner speech decoding was evaluated in sessions t12.2023.11.28, t15.2024.03.03, and t16.2024.04.29 (reported in Fig. 3). Each session began with a “diagnostic” block. This block was used to calculate threshold values and filters for online LRR that would be used for the rest of the session for T12 and T16. For T15, filter and LRR parameters were recalculated after every experimental block. Then, we collected “open-loop” blocks of sentences (3-5 blocks of 40-50 sentences per block) as training data, with no real-time decoder active. We collected each sentence block twice, once with the participant performing inner speech and once with the participant performing attempted speech. We then trained a decoder using the inner speech blocks, as well as using historical attempted speech data that had previously been collected with each participant (see Tables 5-7 below), using decoding methods reported previously in ^1,2^. The architecture of the RNN includes a unique input layer for each day, enabling historical attempted speech data to be successfully added during training without hurting performance on inner speech days. The attempted blocks from the same day were not used to train the online decoder. Next, after the decoder was trained, we collected some additional closed-loop blocks to adjust the rolling mean and z-scoring to account for any nonstationarities that may have accrued during training. For T15 these additional blocks also served as training data, as the model was retraining online as described in ^2,15^. T12 and T16 did not have online retraining. Then we collected a final closed-loop evaluation block of the 50 sentences from ^16^, also used for evaluation in ^1,16^.

### 4.2 Vocabulary and Sentence Selection

To demonstrate a proof-of-concept inner speech neuroprostheses we used a 50-word vocabulary ^1,16^. All of the inner speech sentences were constructed out of this 50-word vocabulary and the evaluation sentence set was taken directly from ^1,16^. There were no overlapping sentences between the training and evaluation sets. The historical attempted data used to supplement the training used sentences from an open vocabulary and were taken from the switchboard corpus as described in ^1^.

### 4.3 Online Decoder Training Data

To assist in training the online decoder for inner speech, we not only used the training blocks of sentences spoken covertly, but also historical attempted or mimed data. Each day of data had a separate input transformation layer that was trained from scratch as described in ^1^. The total number and type of sentences used to train the models which were evaluated in real-time are described below. Note, that since T15’s model employed online retraining, even the sentences decoded in real-time were involved in continually retraining the model throughout the session up to and including during the evaluation block. T12’s and T16’s decoders did not utilize online retraining.

#### 4.3.1 T12

**Table 5:**
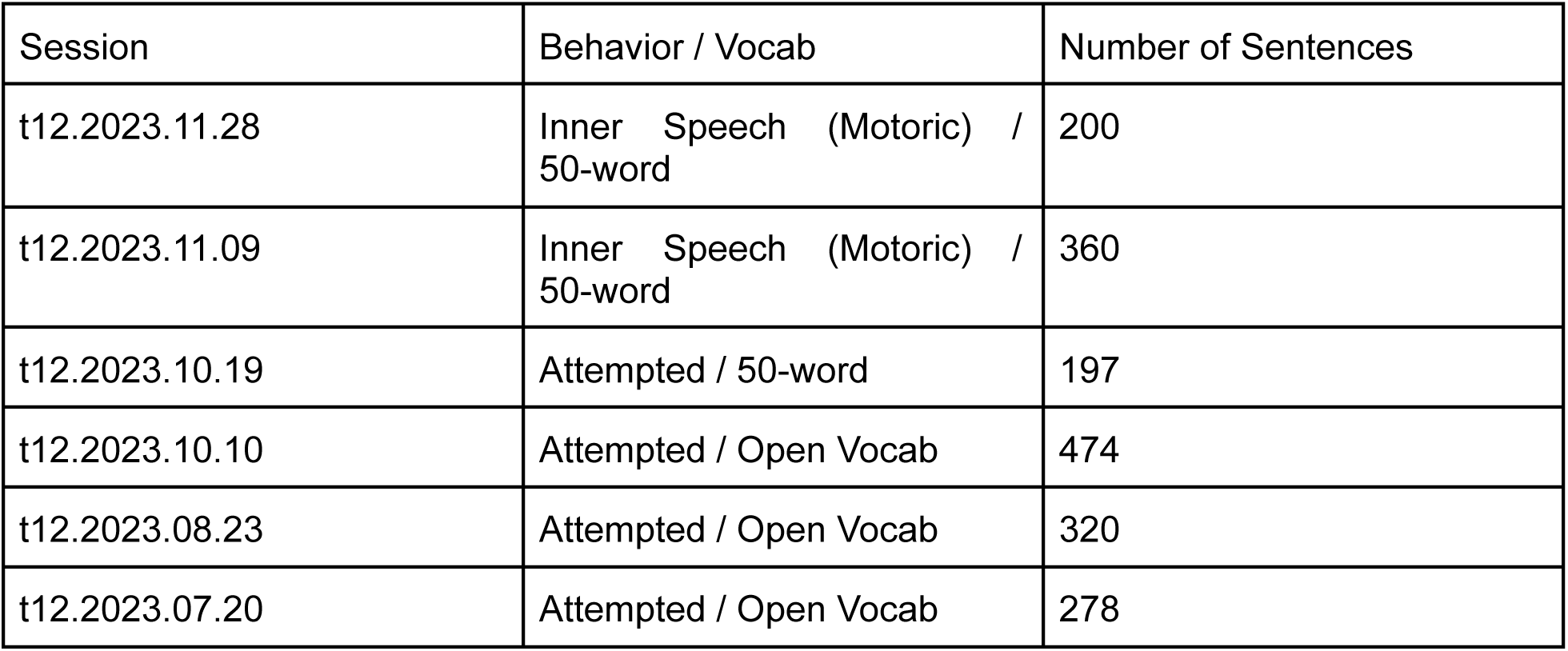
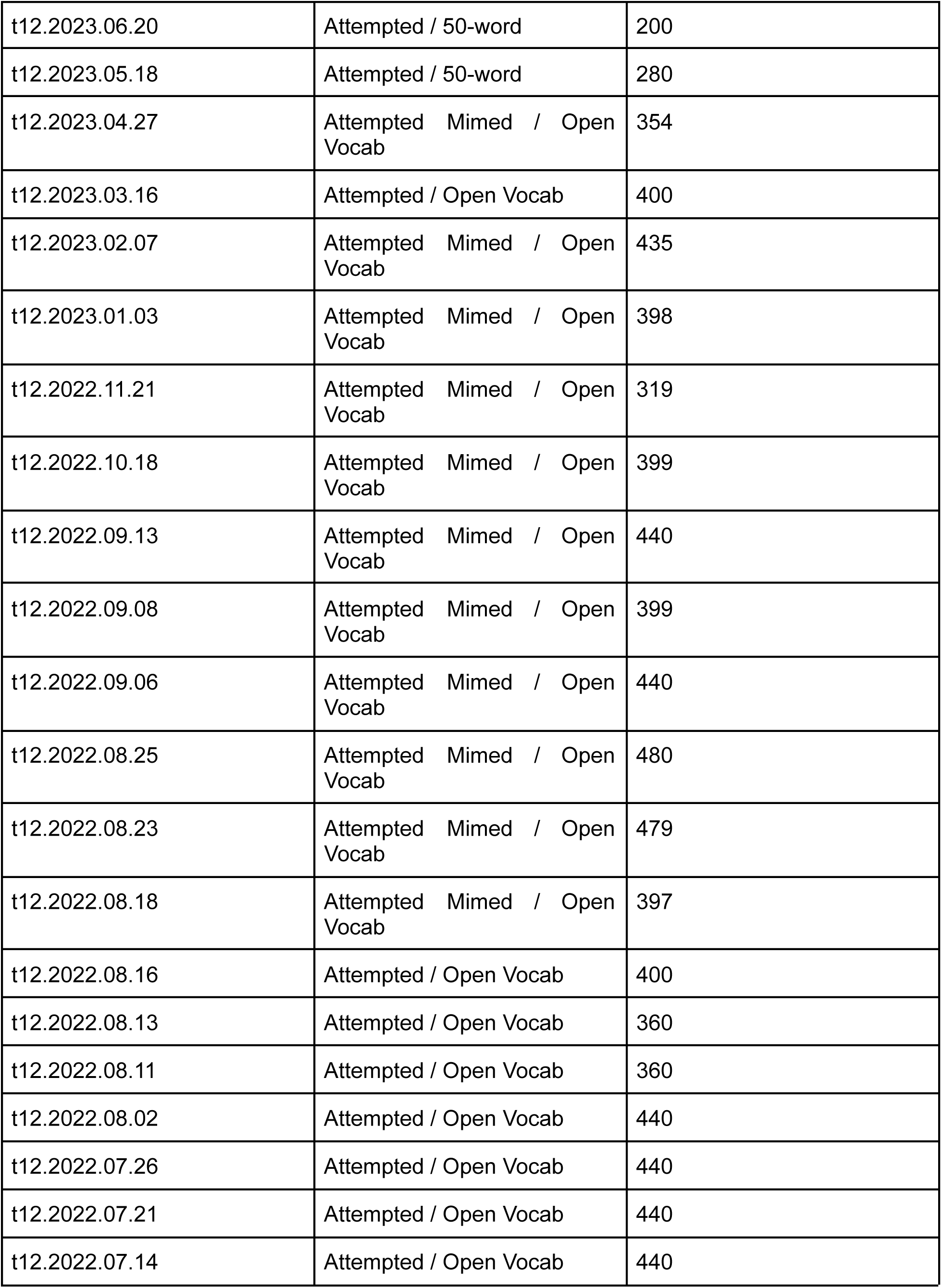
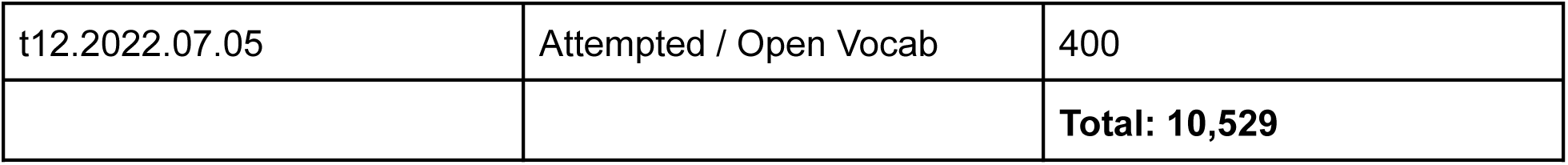
T12 Online Decoder Training Data.

| Session | Behavior / Vocab | Number of Sentences |
| --- | --- | --- |
| t12.2023.11.28 | Inner Speech (Motoric) / 50-word | 200 |
| t12.2023.11.09 | Inner Speech (Motoric) / 50-word | 360 |
| t12.2023.10.19 | Attempted / 50-word | 197 |
| t12.2023.10.10 | Attempted / Open Vocab | 474 |
| t12.2023.08.23 | Attempted / Open Vocab | 320 |
| t12.2023.07.20 | Attempted / Open Vocab | 278 |
| t12.2023.06.20 | Attempted / 50-word | 200 |
| t12.2023.05.18 | Attempted / 50-word | 280 |
| t12.2023.04.27 | Attempted Mimed / Open Vocab | 354 |
| t12.2023.03.16 | Attempted / Open Vocab | 400 |
| t12.2023.02.07 | Attempted Mimed / Open Vocab | 435 |
| t12.2023.01.03 | Attempted Mimed / Open Vocab | 398 |
| t12.2022.11.21 | Attempted Mimed / Open Vocab | 319 |
| t12.2022.10.18 | Attempted Mimed / Open Vocab | 399 |
| t12.2022.09.13 | Attempted Mimed / Open Vocab | 440 |
| t12.2022.09.08 | Attempted Mimed / Open Vocab | 399 |
| t12.2022.09.06 | Attempted Mimed / Open Vocab | 440 |
| t12.2022.08.25 | Attempted Mimed / Open Vocab | 480 |
| t12.2022.08.23 | Attempted Mimed / Open Vocab | 479 |
| t12.2022.08.18 | Attempted Mimed / Open Vocab | 397 |
| t12.2022.08.16 | Attempted / Open Vocab | 400 |
| t12.2022.08.13 | Attempted / Open Vocab | 360 |
| t12.2022.08.11 | Attempted / Open Vocab | 360 |
| t12.2022.08.02 | Attempted / Open Vocab | 440 |
| t12.2022.07.26 | Attempted / Open Vocab | 440 |
| t12.2022.07.21 | Attempted / Open Vocab | 440 |
| t12.2022.07.14 | Attempted / Open Vocab | 440 |
| t12.2022.07.05 | Attempted / Open Vocab | 400 |
|  |  | **Total: 10,529** |

#### 4.3.2 T15

**Table 6:** T15 Online Decoder Training Data.

| Session | Behavior / Vocab | Number of Sentences |
| --- | --- | --- |
| t15.2024.03.03 | Inner Speech (Motoric) / 50-word | 82 (+100 during online retraining in closed-loop) |
| t15.2024.03.01 | Attempted Speech / Open Vocab | 201 |
| t15.2025.02.28 | Attempted Speech / Open Vocab | 253 |
| t15.2024.02.27 | Attempted Speech / Open Vocab | 173 |
| t15.2024.02.25 | Attempted Speech / Open Vocab | 347 |
| t15.2024.02.21 | Attempted Speech / Open Vocab | 298 |
| t15.2024.02.20 | Attempted Speech / Open Vocab | 327 |
| t15.2024.02.18 | Attempted Speech / Open Vocab | 141 |
| t15.2024.02.16 | Attempted Speech / Open Vocab | 99 |
| t15.2024.02.14 | Attempted Speech / Open Vocab | 156 |
| t15.2023.10.01 | Attempted Speech / Open Vocab | 306 |
| t15.2023.09.03 | Attempted Speech / Open Vocab | 244 |
| t15.2023.09.01 | Attempted Speech / Open Vocab | 332 |
| t15.2023.08.13 | Attempted Speech / Open Vocab | 351 |
| t15.2023.08.11 | Attempted Speech / 50-word | 259 |
|  |  | **Total: 3569 (3669)** |

#### 4.3.3 T16

**Table 7:** T16 Online Decoder Training Data.

| Session | Behavior / Vocab | Number of Sentences |
| --- | --- | --- |
| t16.2024.04.29 | Inner Speech (Auditory) / 50-word | 107 |
| t16.2024.04.26 | Attempted Mimed Speech / Open Vocab | 262 |
| t16.2024.04.19 | Attempted Mimed Speech / Open Vocab | 333 |
| t16.2024.04.05 | Attempted Mimed Speech / Open Vocab | 321 |
| t16.2024.03.22 | Attempted Mimed Speech / 50-word Vocab | 138 |
| t16.2024.03.11 | Attempted Mimed Speech / Open Vocab | 184 |
| t16.2024.02.21 | Attempted Mimed Speech / Open Vocab | 176 |
| t16.2024.02.09 | Attempted Speech / Open 50-word Vocab | 181 |
| t16.2024.02.09 | Attempted Mimed Speech / 50-word Vocab | 180 |
| t16.2024.01.26 | Attempted Speech / Open Vocab | 173 |
| t16.2024.01.12 | Attempted Speech / Open Vocab | 167 |
| t16.2024.01.08 | Attempted Speech / Open Vocab | 187 |
|  |  | **Total: 2409** |

### 4.4 RNN

To transform neural activity evoked by inner speech into a time series of phoneme probabilities, we used a 5-layer, stacked gated recurrent unit RNN as described in ^1^. RNN parameters for T12 were determined based on the results of ^1^ and for T15 from ^2^. T16 parameters were based on parameters optimized for attempted speech and matched those of T12.

To train the model without any ground truth labels (given the participants’ inability to produce intelligible speech) we used a connectionist temporal classification (CTC) loss. We also added two types of artificial noise to help regularize the model. For details about training methods see1.

### 4.5 Language Model

To decode word outputs from the real-time phoneme probabilities given by the RNN, we used an n-gram language model (LM) using the 50 word dictionary ^1^). For details about the construction and training, inference and offline optimization of the language model see ^1^ and ^2^.

### 4.6 Word Error Rate

We evaluated decoding performance using word error rate. Word error rate is defined as the edit distance between the decoded sequence of words and the prompt sentence (i.e. the number of insertions , deletions, or substitutions required to make the sequence of words match exactly). Note that the reported error rates are “aggregate” error rates, which are the result of combining across many independent sentences. To combine data across multiple sentences, we summed the number of errors across all sentences and divided this by the total number of words across all sentences (as opposed to computing error rates first for each sentence separately, and then averaging the rates). This helps prevent very short sentences from overly influencing the results. Confidence intervals for error rates were computed via bootstrap resampling over individual trials and then re-computing the aggregate error rates over the resampled distribution (10,000 resamples).

## 5 Inner speech as a mental strategy for serial recall in upper-extremity motor sequence tasks (Figure 4, Supp. Figure 1,2)

### 5.1 Task Descriptions

#### 5.1.1 Upper-extremity motor tasks without mental strategy instructions

Three tasks were designed to variably elicit verbal or non-verbal short-term memory for the execution of an upper-extremity motor task, with the hypothesis that representations of inner speech for cognition could decodable from speech-motor areas. In a suite of instructed-delay tasks (see 1.5) sequences of upper-extremity movement directions were cued and subsequently recalled during the go period. Removal of the visual sequence cue during the go period enforced the use of short-term memory to reproduce the cued movement sequence. All tasks were described to the participants as an upper-extremity motor task without any explicit instruction about what kind of mental strategy to employ. The 3-element arrows task consisted of sequences of three arrows pointing in one of four directions (↑ → ↓ ←) as well as a ‘Do Nothing’ cue. All possible sequences were used, resulting in 65 conditions. A subset of these conditions using only two directions (↑, →) were used for T16 due to session time constraints. T12 was instructed to sequentially move a joystick in the directions of the displayed arrows, returning to center between sequence positions. The single-element arrows task was identical except that only a single movement direction was cued. This task was not done with T16 due to session time constraints. The 3-element lines task was cued with an image of line segments which participants were instructed to reproduce by drawing. The displayed image showed the starting point and three line segments indicating the three target movements (Supp. Figure 2). After an audible go cue, the image was removed after which participants attempted to reproduce the previously displayed image. After a go period, there was a 1.5 second return period to allow T12 to return the pen tip to the start location. Because T12 retained some control of her arm and hand, ground truth joystick and pen tip trajectories were recorded (see 7.1.4). Due to a greater degree of paralysis, T16 was only able to attempt to do hand movements for these tasks.

#### 5.1.2 Upper extremity motor tasks with instructed mental strategy

For the verbal and visual memory upper-extremity motor tasks, the same task design was used but with different instructions for delay and go period mental strategies. Cues consisted of sequences of three arrows pointing either up (↑) or right (→). During the go period the arrows were removed and participants were instructed to attempt to draw a sequence of line segments in the direction of the arrows similar to the 3-element lines. Pen tip trajectories were recorded during drawing (see 5.1.4). For verbal memory tasks, participants were instructed to use inner speech as a mental strategy for short-term memorization of the cued arrow sequence. For visual memory tasks participants were instructed to use visual short-term memory and to suppress any covert speech about arrow directions. Participants were given time to practice until they felt comfortable with the task and reported being able to reliably engage each mental strategy. Ground truth hand movements were recorded as pen-tip trajectories (see 5.1.4) for T12 and are plotted in Supplementary Figure 2.

#### 5.1.3 Analogous overt speech sequence recall task

The previously described verbal memory task (7.1.2) was compared with a speaking task in which T12 was presented with the same direction sequences but via a recorded audio cue. The behavioral instruction was to attempt to speak the directions. Due to limited session time, T16 performed a smaller set of conditions consisting of a single direction (either “up” or “right”) rather than sequences of three directions, and cues were presented as text. For mental strategy T12 was instructed not to change how she would naturally recall and speak the audio-cued direction sequence.

**Table 8:** Summary of hand-motor tasks.

| **Table 8: Summary of hand-motor tasks** |  |  |  |  |  |  |  |  |
| --- | --- | --- | --- | --- | --- | --- | --- | --- |
| **Participant** | **Task Name** | **Instructed Behavior** | **Instructed Mental strategy** | **Number of conditions** | **Repetitions per condition** | **Delay time (s)** | **Go time (s)** | **Return time (s)** |
| T12 | 3-element arrows | Move the joystick in the direction of the arrow cues in sequence | N/A | 65 | 14 | 2.5 | 3 | 4 |
| T12 | single-element arrows | Move the joystick in the direction of the arrow cue | N/A | 5 | 30 | 2.5 | 2.5 | 0 |
| T12 | 3-element lines | Draw three distinct lines to reproduce to image cue | N/A | 17 | 17 | 2.5 | 4 | 1.5 |
| T12 | verbal memory | Draw three distinct lines to reproduce to image cue | Use your inner voice to memorize the cued movement sequence | 17 | 20 | 2.5 | 4 | 1.5 |
| T12 | visual memory | Draw three distinct lines to reproduce to image cue | Use a mental image of the cue to remember what to draw | 17 | 20 | 2.5 | 4 | 1.5 |
| T12 | verbal memory | Draw three distinct lines to reproduce to image cue | Use your inner voice to memorize the cued movement sequence | 9 | 20 | 2.5 | 4 | 1.5 |
| T12 | overt speech | Attempt to speak to reproduce the audio cue of three directions | Prepare your speech as you normally would | 9 | 20 | 3.5 | 4.5 | 0 |
| T16 | 3-element arrows | Attempt to draw in the direction of the arrow cues in sequence. | N/A | 8 | 20 | 2 | 4 | 1 |
| T16 | 3-element lines | Attempt to draw in the direction of the arrow cues in sequence. | N/A | 8 | 20 | 2 | 4 | 1 |
| T16 | visual memory | Attempt to draw in the direction of the arrow cues in sequence. | Use a mental image of the cue to remember what to draw | 8 | 20 | 2 | 4 | 1 |
| T16 | verbal memory | Attempt to draw in the direction of the arrow cues in sequence. | Use your inner voice to memorize the cued movement sequence | 8 | 20 | 2 | 4 | 1 |
| T16 | Overt speech | Attempt to speak the cued word | Prepare your speech as you normally would | 3 | 20 | 2 | 4 | 1 |

#### 5.1.4 Ground truth hand movement tracking

For tasks requiring joystick movement (3-element arrows, single element-arrow), T12’s hand motor activity was recorded using a Logitech Extreme 3D Pro Gaming Joystick. For drawing tasks (3-element lines, inner speech, no inner speech), T12 was instructed to draw on a 15 inch LCD writing tablet (ERUW Shenzhen Lei Rui Technology Co., Ltd) using a stylus. Lines were erased between blocks. The starting position was indicated by drawing an X on the tablet before each block. The Optitrack v120 Trio was used to track the three dimensional positions of the stylus and writing tablet. The stylus was fitted with six infrared reflective markers on a modified Optitrack Hand Rigid Bodies Marker Set which was attached to the back of the stylus using a custom 3d printed mount. Six additional infrared reflective markers were affixed to two adjacent sides of the writing tablet which faced the Optitrack camera using Optitrack Marker Bases in order to estimate the writing plane. Both the stylus and the writing surface were recorded in real time as custom rigid bodies with the three dimensional coordinates of all infrared markers recorded using Motive. The trajectory of the stylus tip on the trackpad was estimated by using the stylus location and orientation as well as the 2d writing surface. At each time sample the stylus rigid body was represented as a quaternion. Tip location was estimated from the recorded quaternion by subtracting the quaternion of a reference stylus with measured tip location. The writing plane was estimated from selecting three points from the tablet markers. Finally, the stylus tip location within the 2 dimensional coordinates of the writing plane was estimated by rotating the stylus quaternion along the writing plane’s normal vector. No ground truth motor activity could be collected with T16 due to a higher degree of paralysis limiting her ability to draw.

### 5.2 Using binary decoding to assess sequence encoding per position

Threshold crossing rates were summed over a 2 second window before the go cue to isolate preparatory activity, generating a 64 length vector for each trial and microelectrode array. Decodability of individual positions was estimated by comparing two sequences that differ in only a single position. For T16, a 0.75 second window immediately following the go cue was used. Binary LDA decoders ^17^ were fit for each pair of sequences that differed in only one position. Five fold cross validation was used to prevent overfitting. Boxplots for each sequence position depict the distribution of decoding accuracies across all possible pairs differing at that position (Figure 4).

### 5.3 Confidence interval estimation via bootstrap

Confidence intervals for decoding performance at each sequence position were calculated by resampling decoder predictions. For each decoder, the cross-validated predictions were resampled with replacement. Resampled predictions from all decoders for a specific sequence position were joined to estimate the per-position decoding accuracy. This was repeated 10000 times to estimate the distribution of per-position decoding accuracy. Significance was assessed by taking the 0.025th quantile and comparing it to the null hypothesis of chance-level decoding accuracy (0.5).

#### 5.3.1 Paired increase in decoding accuracy for instructed verbal vs visual memory task

To assess the significance of the effect of explicit instruction to use either verbal or visual memory for sequence recall, the distribution of increases in decoding accuracy for paired decoders differing only in instructed mental strategy was estimated per position. For each decoder, the cross validated predictions were resampled with replacement to estimate per-decoder resampled accuracy. Then, paired decoding accuracies for data only differing in instructed mental strategy were subtracted in order to compute the increase in decoding accuracy due to instruction to use a verbal versus visual mental strategy. Finally, this was repeated 10000 times to estimate the distribution of increased decoding accuracy for each position. Significance was assessed by taking the 0.025th quantile and comparing it to the null hypothesis of no increase in decoding accuracy.

### 5.4 Cross task positional decoding

To assess whether sequence representation in the hand-motor task was indeed due to inner speech, we tested whether decoders trained on a speech sequence task could generalize to the hand-motor task when inner speech is used. LDA decoders were fit similarly to 5.2 except that train and test data were from different tasks. No cross validation was used as there was no potential for overfitting.

## 6 Relationship of overt to covert speech (Figure 5)

While the ‘isolated’ verbal behavior experiments (section 3) allowed us to explore a large number of behaviors, it did not allow us to assess whether mean firing rates may change between behaviors, because any firing rate differences between blocks could also be caused by spurious neural non-stationarity (firing rate drift). In order to assess whether large mean firing rate differences exist between overt and covert speech (which could be a useful cue for a decoder to distinguish between them), we ran a follow-up task in which overt, covert and listening conditions were randomly interleaved within an experimental block. When trials are interleaved within a block, any differences in mean between behaviors is preserved when performing block-wise mean subtraction to remove firing rate drift across time. .

### 6.1 Interleaved Verbal Behavior Instructed-Delay Task

This task included three behaviors randomly interleaved within the same experimental block (attempted speech, motoric inner speech, and listening for T12; attempted speech, imagined listening and listening for T15; attempted mimed speech, auditory inner speech, and listening for T16).

### 6.2 Cross-validated Euclidean neural distance within and across behaviors (Figure 5D)

In Figure 5D, we estimated the average Euclidean distance in neural state space between all 21 word pairs within a behavior (“within covert” and “within overt” bars) and between all 7 matching word pairs across behaviors (“Overtness Dim.” bars) following the methods described in ^14^. Across-behavior distances estimate the size of the change in mean firing rates between the behaviors (which together constitute an “overtness dimension” in neural state space), while within-behavior distances estimate the size of the neural modulation evoked by words. Confidence intervals (95%) were estimated for the mean within-behavior distances (21 points) and across-behavior distances (7 points) by assuming the points were normally distributed.

## 7 Offline continuous inner speech analysis (Figure 3E, Figure 6)

### 7.1 Overt and Covert Training and Test Sets

To assess decoding performance for continuous overt and covert speech, we collected data where participants spoke the same sentences both overtly and covertly, and evaluated decoders offline using train / test splits of those sentences that were identical across behaviors. All sentences were constructed from the 50-word vocabulary. If a sentence was removed due to an interruption during data collection, the corresponding sentence in the other behavior was also removed. Interruptions during data collection that warranted trial removal consisted of coughing bouts, participant care needs, interruption by person in the room, or loud noises that masked the sound of the task computer, (i.e. from a passing train), all of which we believe prevented the participant from being able to fully perform the task during the given trial.

**Table 9:** Overt and Covert Training and Test Sets By Participant.

| Participant | Covert Behavior | Overt Behavior | # Sentences Train (per behavior) | # Sentences Test (per behavior) |
| --- | --- | --- | --- | --- |
| T12 | Inner Speech (Motoric) | Attempted Speech | 130 | 30 |
| T15 | Inner Speech<br>(Motoric) | Attempted<br>Speech | 164 | 30 |
| T16 | Inner Speech<br>(Auditory) | Attempted<br>Mimed Speech | 85 | 30 |

### 7.2 Offline decoding evaluation (Figure 3E)

To compare decoding performance directly between overt speech (attempted speech for T12 and T15; attempted mimed for T16) and inner speech (Figure 3E), we trained offline models using only the continuous 50-word sentences collected on the inner speech decoding evaluation day for each participant (Table 9 above). We trained 10 seeds each of models trained on the overt condition and the covert condition training sets, and evaluated on each behavior’s corresponding test set, yielding directly comparable performance numbers for covert and overt speech. The test set consisted of the final 30 sentences collected for each participant.

To evaluate cross-decoding performance (i.e., how well a decoder trained on overt speech could inadvertently decode covert speech), we simply evaluated the overt models on the covert test set. RNN hyperparameters for all offline analyses were taken from our prior work ^1,2^.

To estimate a chance level word error rate that could be expected if the RNN output random sentences unrelated to the prompt, decoded sentences were shuffled relative to the ground truth sentences. We did this 1,000 times for each of the 10 seeds of each scenario investigated (trained and tested on overt only, trained and tested on covert only, and trained on overt but tested on covert). All actual word error rate confidence intervals were better than the determined chance level for each scenario (for each bar in Figure 3E).

### 7.3 “Imagery silenced” performance evaluation (Figures 6C-F)

#### 7.3.1 Decoding performance by training strategy (Figure 6C-D)

We evaluated the offline word error rate of “imagery-silenced” and “imagery-naive” training strategies, when tested on overt speech (Figure 6C), following the methods described above in section 7.2. Separately, we evaluated how often decoders trained with each strategy erroneously produced output on covert speech trials (Figure 6D). If any token other than “>SIL<” was produced by the RNN, the trial was considered a failure.

#### 7.3.2 Correlation of logits (Figure 6E)

To further examine the effect of the "imagery-silenced" training strategy on decoder output, we examined the phoneme probabilities (logits) given by the RNN decoder for paired covert and overt trials. We reasoned that in the imagery-naive case, the decoder might inadvertently output similar probabilities for matched covert and overt trials, given the high correlation between overt and covert speech. In the imagery-silenced case, however, outputs should ideally be much less similar. We quantified this by first time-warping the decoder outputs to align matched overt and covert trials (thereby removing natural variation in speaking rate), and then computing the correlation (Pearson r) between time warped logit time series.

##### 7.3.3.1 Dynamic Time Warping (DTW)

In order to align the covert and overt trials (which might have similar content but be misaligned in time due to different rates of speech and typical behavioral variation across trials), we used dynamic time warping ^18^. To do this, we used the python dynamic time warping package ^18^ with a ‘symmetric2’ step pattern and slanted band window of a 100ms size. This allows for some flexible time alignment while also constraining variation so that time-warping cannot be too extreme and overfit to noise.

##### 7.3.3.2 Correlations

After aligning corresponding covert and overt logit time series with Dynamic Time Warping, we then calculated the correlation (Pearson’s r) for each sentence separately. For each sentence, a correlation was computed for each of the 39 phoneme logits, and correlations were averaged to yield a single value. This procedure was done for all 30 test-set sentences and all 10 decoder seeds, for each training strategy. Bar heights in Figure 6E represent the average across all 30 sentences and 10 seeds. Chance levels were computed by shuffling sentences within each behavior, such that sentences were no longer paired when correlating. One thousand shufflings were performed for each of 10 seeds; the mean of this distribution was taken as the chance value (dashed line). Confidence intervals (95%) were computed via bootstrap resampling over individual trials and then re-computing the average correlations over the resampled distribution (10,000 resamples).

## 8 Statistics

| Result | Statistical Details |
| --- | --- |
| Fig 1D | 95% confidence intervals of the mean were computed by assuming that the data are normally distributed. |
| Fig 1E | 95% confidence intervals for decoding accuracy were computed using a Clopper-Pearson binomial confidence interval, computed over all independent test trials concatenated across the 10 folds. Accuracy was deemed to be statistically significantly above chance if the lower bound of the |
|  | confidence interval was greater than the chance value of 14.3%. |
| Figure 3D,3E, 6C | Confidence intervals for error rates were computed via bootstrap resampling over individual trials and then re-computing the aggregate error rates over the resampled distribution (10,000 resamples) |
| Figure 4BDF, Supplementary Figure 1BDF | Whiskers depict 1.5 times the interquartile range. Box depicts the 25th and 75th quantiles and the median. For B and F, significance was assessed by computing the 95% confidence interval of decoding accuracy per sequence position via bootstrap and comparing it to the null hypothesis of chance performance (see 5.3). For D, significance was assessed by computing the 95% confidence interval of decoder performance increase due to mental strategy instruction and comparing it to the null hypothesis of no change in decoder performance (see 5.3.1). |
| Fig 5D | 95% confidence intervals of the mean euclidean distance between conditions were computed by assuming that the data are normally distributed. |
| Figure 6D | Bar heights indicate the percentage of “failed” trials (trials where words were decoded) out of all covert test trials. 95% confidence intervals were calculated in a similar bootstrap manner as described in Section 5 for WER in which 1,000 resamplings were taken for each of the 10 model seeds These 10,000 samples were used to compute the confidence interval (2.5 to 97.5 percentiles). |
| Figure 6E | To calculate chance levels (dashed lines in 6E), , sentences were shuffled within each behavior, so that correlations were no longer taken over matching sentences.We did this 1,000 times for each of the 10 model seeds. We took the mean of these 10,000 values as the chance level. |
|  | Confidence intervals (95%) were computed via bootstrap resampling over individual trials and then re-computing the average correlations over the resampled distribution (10,000 resamples). |

## Supplement

**Supplementary Figure 1:**
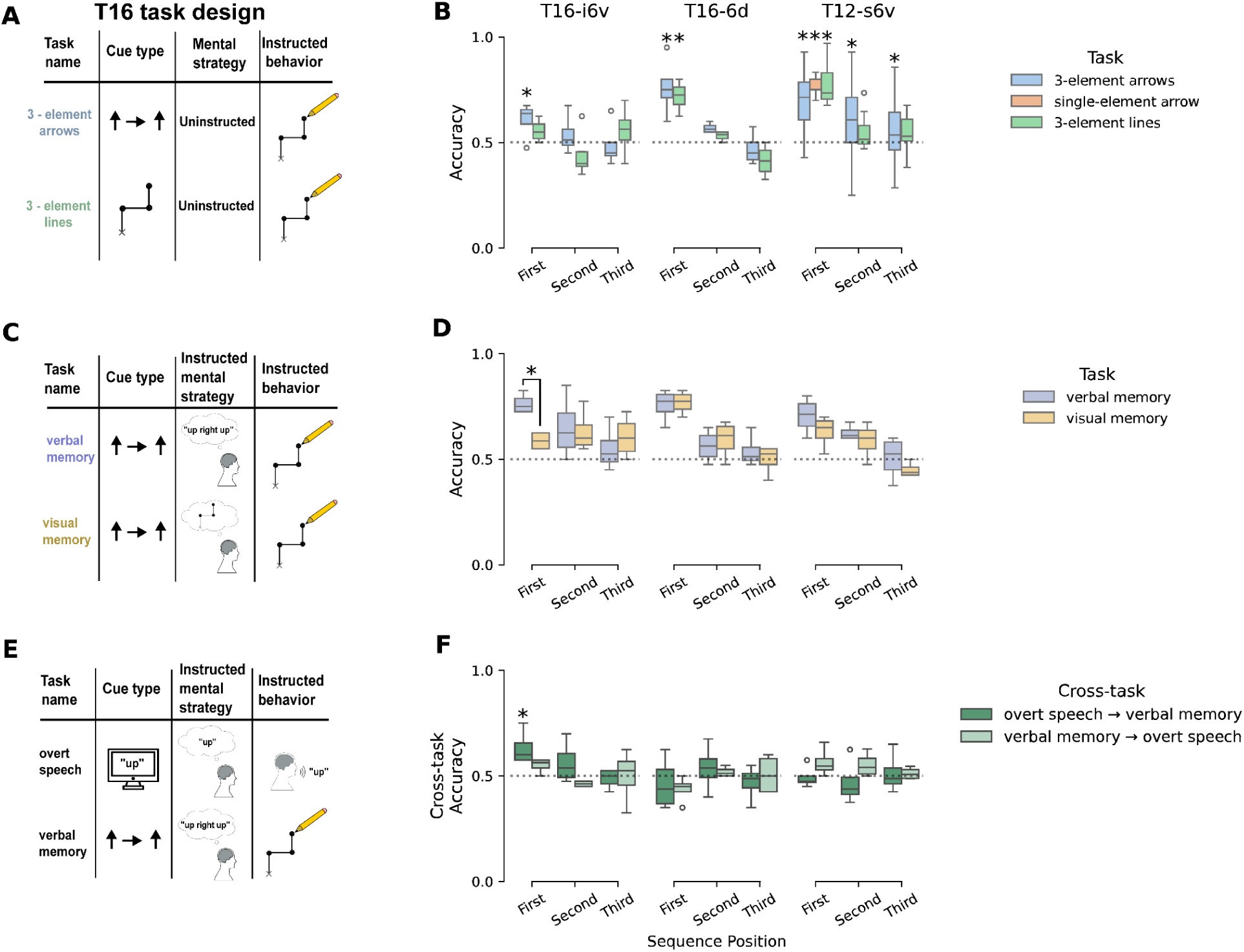
Inner speech for task execution is also decodable in i6v in T16 whereas areas 6d in T16 and s6v in T12 exhibit hand - motor tuning. A) Participant T16 also completed the 3-element arrows and lines tasks without explicit instruction for mental strategy. Due to T16’s upper-extremity paralysis, attempted drawing was instructed as the desired behavior for sequence recall rather than actual drawing as was instructed for T12. Task design for T12 is describe in Figure 4A. B) Decodability of sequence position was assessed as in Figure 4. For T16, a window of neural activity from the first 1.5 seconds after the go cue was used to fit decoders. For area i6v in T16, only the 3-element arrow task elicited significant neural representation of the first sequence position. C) Participant T16 performed two versions of the three-element arrows task, but with explicit instruction to either use or suppress inner speech for short-term memory of the arrow sequence. D) Same as B but for tasks that only differed in instructed mental strategy. Instruction to use verbal mental strategy significantly increased decoding accuracy of the first position in area i6v in T16 (mean decoding accuracy 0.61, 95% CI 0.53-0.68) but not in areas 6d in T16 nor area s6v in T12. E) T16 was visually cued by text to speak a direction to test whether the covert for short term memory in i6v had a shared representation with overt speech. F) Same as B except decoders are trained on overt (covert) speech and tested on covert (overt) speech. For T16 i6v, only overt → covert decoders could generalize above chance for the first position. Decoders did not generalize well for area 6d in T16.

**Supplementary Figure 2:**
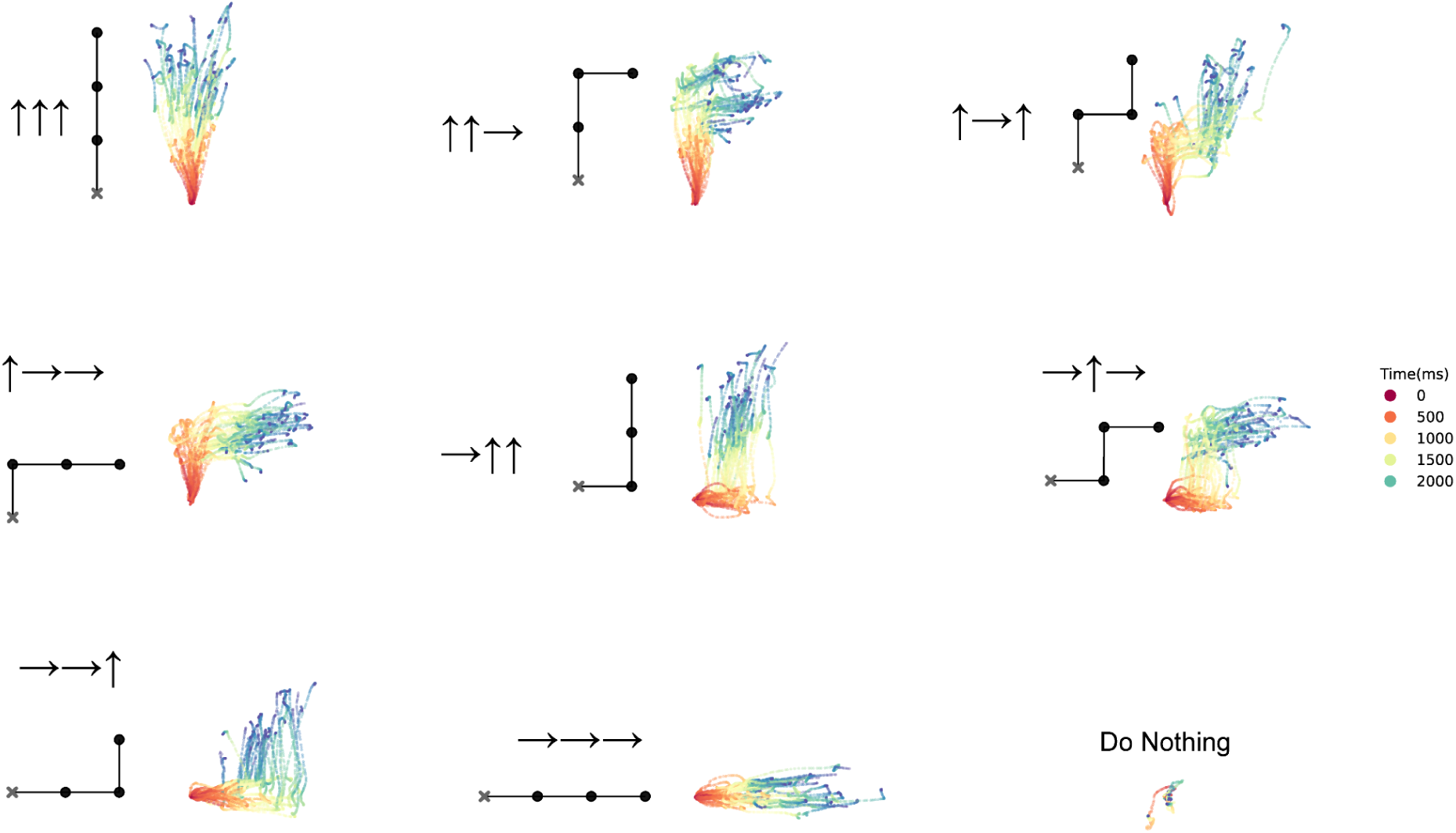
Error-free and consistent drawing behavior by T12. T12 has limited control of upper extremities, allowing ground truth recording of drawing behavior during upper extremity sequential motor tasks described in Figure 4C. Each movement sequence was cued either with arrows or or lines indicating the target drawn movement sequence as well as a ’Do Nothing’ condition in which T12 attempted to not move. T12 performed the behavior with 100% accuracy as judged by manual annotation of pen tip trajectories from individual trials.

## References

1. Willett, F. R. et al. A high-performance speech neuroprosthesis. Nature 620, 1031–1036 (2023).

2. Metzger, S. L. et al. A high-performance neuroprosthesis for speech decoding and avatar control. Nature 620, 1037–1046 (2023).

3. Card, N. S. et al. An accurate and rapidly calibrating speech neuroprosthesis. N. Engl. J. Med. 391, 609–618 (2024).

4. Soldado-Magraner, J. et al. Applying the IEEE BRAIN neuroethics framework to intra-cortical brain-computer interfaces. J. Neural Eng. 21, (2024).

5. Brown, C. M. L. Neurorights, Mental Privacy, and Mind Reading. Neuroethics 17, 34 (2024).

6. van Stuijvenberg, O. C., Samlal, D. P. S., Vansteensel, M. J., Broekman, M. L. D. & Jongsma, K. R. The ethical significance of user-control in AI-driven speech-BCIs: a narrative review. Front. Hum. Neurosci. 18, (2024).

7. Pels, E. G. M., Aarnoutse, E. J., Ramsey, N. F. & Vansteensel, M. J. Estimated Prevalence of the Target Population for Brain-Computer Interface Neurotechnology in the Netherlands. Neurorehabil. Neural Repair 31, 677–685 (2017).

8. Pandarinath, C. et al. High performance communication by people with paralysis using an intracortical brain-computer interface. Elife 6, (2017).

9. Gilja, V. et al. Clinical translation of a high-performance neural prosthesis. Nat. Med. 21, 1142–1145 (2015).

10. Hochberg, L. R. et al. Neuronal ensemble control of prosthetic devices by a human with tetraplegia. Nature 442, 164–171 (2006).

11. Schalk, G. et al. Two-dimensional movement control using electrocorticographic signals in humans. J. Neural Eng. 5, 75–84 (2008).

12. Nuyujukian, P. et al. Cortical control of a tablet computer by people with paralysis. PLoS One 13, e0204566 (2018).

13. Simeral, J. D., Kim, S. P., Black, M. J., Donoghue, J. P. & Hochberg, L. R. Neural control of cursor trajectory and click by a human with tetraplegia 1000 days after implant of an intracortical microelectrode array. J. Neural Eng. 8, 025027 (2011).

14. Hochberg, L. R. et al. Reach and grasp by people with tetraplegia using a neurally controlled robotic arm. Nature 485, 372–375 (2012).

15. Collinger, J. L. et al. High-performance neuroprosthetic control by an individual with tetraplegia. Lancet 381, 557–564 (2013).

16. Ajiboye, A. B. et al. Restoration of reaching and grasping movements through brain-controlled muscle stimulation in a person with tetraplegia: a proof-of-concept demonstration. Lancet 389, 1821–1830 (2017).

17. Willett, F. R., Avansino, D. T., Hochberg, L. R., Henderson, J. M. & Shenoy, K. V. High-performance brain-to-text communication via handwriting. Nature 593, 249–254 (2021).

18. Moses, D. A. et al. Neuroprosthesis for Decoding Speech in a Paralyzed Person with Anarthria. N. Engl. J. Med. 385, 217–227 (2021).

19. Anumanchipalli, G. K., Chartier, J. & Chang, E. F. Speech synthesis from neural decoding of spoken sentences. Nature 568, 493–498 (2019).

20. Silva, A. B., Littlejohn, K. T., Liu, J. R., Moses, D. A. & Chang, E. F. The speech neuroprosthesis. Nat. Rev. Neurosci. 25, 473–492 (2024).

21. Baars, B. How Brain Reveals Mind Neural Studies Support the Fundamental Role of Conscious Experience. Journal of Consciousness Studies 10, 100–114 (2003).

22. Hurlburt, R. T., Heavey, C. L. & Kelsey, J. M. Toward a phenomenology of inner speaking. Conscious. Cogn. 22, 1477–1494 (2013).

23. D’Argembeau, A., Renaud, O. & Van der Linden, M. Frequency, characteristics and functions of future-oriented thoughts in daily life. Appl. Cogn. Psychol. 25, 96–103 (2011).

24. Hubbard, T. L. Auditory imagery: empirical findings. Psychol. Bull. 136, 302–329 (2010).

25. Alderson-Day, B. & Fernyhough, C. Inner Speech: Development, Cognitive Functions, Phenomenology, and Neurobiology. Psychol. Bull. 141, 931–965 (2015).

26. Baddeley, A. Working Memory. Science 255, 556–559 (1992).

27. Baddeley, A., Chincotta, D. & Adlam, A. Working memory and the control of action: evidence from task switching. J. Exp. Psychol. Gen. 130, 641–657 (2001).

28. Sokolov, E. N., Spinks, J. A., Näätänen, R. & Lyytinen, H. The orienting response in information processing. 368, (2002).

29. Newton, A. M. & de Villiers, J. G. Thinking while talking: adults fail nonverbal false-belief reasoning. Psychol. Sci. 18, 574–579 (2007).

30. Hardy, J. Speaking clearly: A critical review of the self-talk literature. Psychol. Sport Exerc. 7, 81–97 (2006).

31. Dolcos, S. & Albarracin, D. The inner speech of behavioral regulation: Intentions and task performance strengthen when you talk to yourself as a You. Eur. J. Soc. Psychol. 44, 636–642 (2014).

32. Perrone-Bertolotti, M., Rapin, L., Lachaux, J. P., Baciu, M. & Lœvenbruck, H. What is that little voice inside my head? Inner speech phenomenology, its role in cognitive performance, and its relation to self-monitoring. Behav. Brain Res. 261, 220–239 (2014).

33. Baddeley, A., Eldridge, M. & Lewis, V. The role of subvocalisation in reading. The Quarterly Journal of Experimental Psychology Section A 33, 439–454 (1981).

34. Soroush, P. Z. et al. The nested hierarchy of overt, mouthed, and imagined speech activity evident in intracranial recordings. Neuroimage 269, 119913 (2023).

35. Tankus, A., Solomon, L., Aharony, Y., Faust-Socher, A. & Strauss, I. Machine learning algorithm for decoding multiple subthalamic spike trains for speech brain-machine interfaces. J. Neural Eng. 18, 066021 (2021).

36. Proix, T. et al. Imagined speech can be decoded from low- and cross-frequency intracranial EEG features. Nat. Commun. 13, 48 (2022).

37. Bookheimer, S. Y., Zeffiro, T. A., Blaxton, T., Gaillard, W. & Theodore, W. Regional cerebral blood flow during object naming and word reading. Hum. Brain Mapp. 3, 93–106 (1995).

38. Rosen, H. J. et al. Neural correlates of recovery from aphasia after damage to left inferior frontal cortex. Neurology 55, 1883–1894 (2000).

39. Palmer, E. D. et al. An event-related fMRI study of overt and covert word stem completion. Neuroimage 14, 182–193 (2001).

40. Huang, J., Carr, T. H. & Cao, Y. Comparing cortical activations for silent and overt speech using event-related fMRI. Hum. Brain Mapp. 15, 39–53 (2002).

41. Shuster, L. I. & Lemieux, S. K. An fMRI investigation of covertly and overtly produced mono- and multisyllabic words. Brain Lang. 93, 20–31 (2005).

42. Frings, M. et al. Cerebellar involvement in verb generation: an fMRI study. Neurosci. Lett. 409, 19–23 (2006).

43. Basho, S., Palmer, E. D., Rubio, M. A., Wulfeck, B. & Müller, R.-A. Effects of generation mode in fMRI adaptations of semantic fluency: paced production and overt speech. Neuropsychologia 45, 1697–1706 (2007).

44. Forn, C. et al. A comparison of brain activation patterns during covert and overt paced auditory serial addition test tasks. Hum. Brain Mapp. 29, 644–650 (2008).

45. Kielar, A., Milman, L., Bonakdarpour, B. & Thompson, C. K. Neural correlates of covert and overt production of tense and agreement morphology: Evidence from fMRI. J. Neurolinguistics 24, 183–201 (2011).

46. Pei, X. et al. Spatiotemporal dynamics of electrocorticographic high gamma activity during overt and covert word repetition. Neuroimage 54, 2960–2972 (2011).

47. de Borman, A. et al. Imagined speech event detection from electrocorticography and its transfer between speech modes and subjects. *Commun*. Biol. 7, 818 (2024).

48. Martin, S., Iturrate, I., Millán, J. D. R., Knight, R. T. & Pasley, B. N. Decoding Inner Speech Using Electrocorticography: Progress and Challenges Toward a Speech Prosthesis. Front. Neurosci. 12, 422 (2018).

49. Zhang, W., Liu, Y., Wang, X. & Tian, X. The dynamic and task-dependent representational transformation between the motor and sensory systems during speech production. Cogn. Neurosci. 11, 194–204 (2020).

50. Ikeda, S. et al. Neural decoding of single vowels during covert articulation using electrocorticography. Front. Hum. Neurosci. 8, 125 (2014).

51. Martin, S. et al. Corrigendum: Word pair classification during imagined speech using direct brain recordings. Sci. Rep. 7, 44509 (2017).

52. Angrick, M. et al. Real-time synthesis of imagined speech processes from minimally invasive recordings of neural activity. Commun Biol 4, 1055 (2021).

53. Wandelt, S. K. et al. Representation of internal speech by single neurons in human supramarginal gyrus. Nat Hum Behav 8, 1136–1149 (2024).

54. Glasser, M. F. et al. A multi-modal parcellation of human cerebral cortex. Nature 536, 171–178 (8/2016).

55. Druckmann, S. & Chklovskii, D. Over-complete representations on recurrent neural networks can support persistent percepts. Adv. Neural Inf. Process. Syst. 541–549 (2010).

56. Druckmann, S. & Chklovskii, D. B. Neuronal circuits underlying persistent representations despite time varying activity. Curr. Biol. 22, 2095–2103 (2012).

57. Vyas, S., Golub, M. D., Sussillo, D. & Shenoy, K. V. Computation Through Neural Population Dynamics. Annu. Rev. Neurosci. 43, 249–275 (2020).

58. Kaufman, M. T., Churchland, M. M., Ryu, S. I. & Shenoy, K. V. Cortical activity in the null space: permitting preparation without movement. Nat. Neurosci. 17, 440–448 (2014).

59. Li, N., Daie, K., Svoboda, K. & Druckmann, S. Robust neuronal dynamics in premotor cortex during motor planning. Nature 532, 459–464 (2016).

60. Duque, J. & Ivry, R. B. Role of corticospinal suppression during motor preparation. Cereb. Cortex 19, 2013–2024 (2009).

61. Miyake, A., Emerson, M. J., Padilla, F. & Ahn, J.-C. Inner speech as a retrieval aid for task goals: the effects of cue type and articulatory suppression in the random task cuing paradigm. Acta Psychol. 115, 123–142 (02/2004).

62. Gilhooly, K. J., Logie, R. H., Wetherick, N. E. & Wynn, V. Working memory and strategies in syllogistic-reasoning tasks. Mem. Cognit. 21, 115–124 (1993).

63. Jarosiewicz, B. et al. Virtual typing by people with tetraplegia using a self-calibrating intracortical brain-computer interface. Sci. Transl. Med. 7, 313ra179 (2015).

64. Perge, J. A. et al. Intra-day signal instabilities affect decoding performance in an intracortical neural interface system. J. Neural Eng. 10, 036004 (2013).

65. Wernicke, C. Der aphasische Symptomencomplex. (1874).

66. Lichtheim, L. & Others. On aphasia. in Broca’s Region 318–347 (Oxford University Press Oxford, 1885).

67. Damasio, A. R. & Geschwind, N. The neural basis of language. Annu. Rev. Neurosci. 7, 127–147 (1984).

68. Wind, J., Chiarelli, B., Bichakjian, B., Nocentini, A. & Jonker, A. Language Origin: A Multidisciplinary Approach. (Springer Science & Business Media, 2013).

69. Pulvermüller, F. Words in the brain’s language. Behav. Brain Sci. 22, 253–79; discussion 280–336 (1999).

70. Braitenberg, V. & Pulvermüller, F. Entwurf einer neurologischen Theorie der Sprache. Sci. Nat. 79, 103–117 (1992).

71. Schippers, A., Vansteensel, M. J., Freudenburg, Z. V. & Ramsey, N. F. Don’t put words in my mouth: Speech perception can generate False Positive activation of a speech BCI. medRxiv (2024) doi:10.1101/2024.01.21.23300437.

72. Dekleva, B. M. et al. Motor cortex retains and reorients neural dynamics during motor imagery. Nat Hum Behav 8, 729–742 (2024).

73. Fedorenko, E., Piantadosi, S. T. & Gibson, E. A. F. Language is primarily a tool for communication rather than thought. Nature 630, 575–586 (2024).

74. Kompa, N. A. Inner speech and ‘pure’ Thought – do we think in language? Rev. Philos. Psychol. 15, 645–662 (2024).

75. Muñoz, J. M. et al. Effects of the first successful lawsuit against a consumer neurotechnology company for violating brain data privacy. Nat. Biotechnol. 42, 1015–1016 (2024).

76. King, B. J., Read, G. J. M. & Salmon, P. M. The risks associated with the use of brain-computer interfaces: A systematic review. Int. J. Hum. Comput. Interact. 1–18 (2022).

## References

1. Willett, F. R. et al. A high-performance speech neuroprosthesis. Nature 620, 1031–1036 (2023).

2. Card, N. S. et al. An accurate and rapidly calibrating speech neuroprosthesis. N. Engl. J. Med. 391, 609–618 (2024).

3. Glasser, M. F. et al. A multi-modal parcellation of human cerebral cortex. Nature 536, 171–178 (2016).

4. Deo, D. R. et al. A mosaic of whole-body representations in human motor cortex. bioRxivorg (2024) doi:10.1101/2024.09.14.613041.

5. Masse, N. Y. et al. Non-causal spike filtering improves decoding of movement intention for intracortical BCIs. J. Neurosci. Methods 236, 58–67 (2014).

6. Young, D. et al. Signal processing methods for reducing artifacts in microelectrode brain recordings caused by functional electrical stimulation. J. Neural Eng. 15, 026014 (2018).

7. Trautmann, E. M. et al. Accurate Estimation of Neural Population Dynamics without Spike Sorting. Neuron 103, 292–308.e4 (2019).

8. Chestek, C. A. et al. Long-term stability of neural prosthetic control signals from silicon cortical arrays in rhesus macaque motor cortex. J. Neural Eng. 8, 045005 (2011).

9. Christie, B. P. et al. Comparison of spike sorting and thresholding of voltage waveforms for intracortical brain-machine interface performance. J. Neural Eng. 12, 016009 (2015).

10. Jarosiewicz, B. et al. Virtual typing by people with tetraplegia using a self-calibrating intracortical brain-computer interface. Sci. Transl. Med. 7, 313ra179 (2015).

11. Perge, J. A. et al. Intra-day signal instabilities affect decoding performance in an intracortical neural interface system. J. Neural Eng. 10, 036004 (2013).

12. Brainard, D. H. The Psychophysics Toolbox. Spat. Vis. 10, 433–436 (1997).

13. Ali, Y. H. et al. BRAND: a platform for closed-loop experiments with deep network models. J. Neural Eng. 21, (2024).

14. Willett, F. R. et al. Hand Knob Area of Premotor Cortex Represents the Whole Body in a Compositional Way. Cell 181, 396–409.e26 (2020).

15. Fan, C. et al. Plug-and-Play Stability for Intracortical Brain-Computer Interfaces: A One-Year Demonstration of Seamless Brain-to-Text Communication. Adv. Neural Inf. Process. Syst. 36, 42258–42270 (2023).

16. Moses, D. A. et al. Neuroprosthesis for Decoding Speech in a Paralyzed Person with Anarthria. N. Engl. J. Med. 385, 217–227 (2021).

17. Pedregosa, F. et al. Scikit-learn: Machine Learning in Python. MACHINE LEARNING IN PYTHON 6 (2011).

18. Giorgino, T. Computing and Visualizing Dynamic Time Warping Alignments in R: The dtw Package. J. Stat. Softw. 31, 1–24 (2009).

